# From tissue to subcellular level : imaging human precision-cut lung slices (PCLS) to gain insight into pandemic bacterial or viral infections

**DOI:** 10.1101/2024.09.04.611185

**Authors:** Sébastien Eymieux, Anne Bull-Maurer, Julien Pichon, Damien Sizaret, Marianne Maquart, Florence Carreras, Maïa Saint-Vanne, Emilie Doz-Deblauwe, Badreddine Bounab, Béatrice Lipan, Lynda Handala, Florentine Chesnel, Julien Burlaud-Gaillard, Fabrizio Mammano, Denys Brand, Antoine Legras, Nathalie Winter, Aude Remot

## Abstract

We describe a method for the generation and deep imaging of human precision-cut lung slices (PCLS). PCLS bridge the gap between *in vivo* and *in vitro* studies, providing a robust system for visualizing events from tissue to subcellular levels in the three-dimensional lung environment, with the preservation of all resident cell types and cell-cell interactions. They also constitute a validated model for studying host cell-pathogen interactions. Here, we detail the generation of human PCLS, followed by their infection and imaging by laser scanning confocal microscopy and transmission electron microscopy (TEM). We establish the conditions for *ex vivo* infection and replication of two pathogens of relevance to human respiratory health: a virus (SARS-CoV-2) and a bacterium (*Mycobacterium tuberculosis,* Mtb). PCLS can be obtained in a single day, infected the next day, and were successfully cultivated for up to a week in this study. Imaging was performed on fixed samples. The preparation of PCLS took one day for confocal imaging and five days for TEM imaging. All procedures are readily adaptable to explore other pathogens and other species and are easy to implement by users with experience in tissue culture. Some specialist equipment (an Alabama tissue slicer) is required for PCLS generation.

## Introduction

Recent advances in three-dimensional (3D) models of the respiratory system have helped improve our understanding of human respiratory biology and pathology (1). These models range from simple submerged systems to complex, perfused constructs, each with its own distinct advantages and limitations. Submerged human airway epithelial cells, for example, provide a straightforward and quick setup for studies of primary cells, but lack the cellular polarization essential to mimic the ciliated and secretory functions observed *in vivo*. By contrast, human airway epithelial cells cultured in air-liquid interface conditions better represent the mucociliary pseudostratified epithelium and can be used to study exposure to airborne substances (2,3). However, this model requires careful handling due to the use of Transwell® systems coated with collagen. Perfusion-based respiratory system models provide better culture conditions, with a constant supply of nutrients, rendering them suitable for long-term studies (4). However, they require specialist equipment and pose a higher risk of contamination. Lung organoids, with their self-organizing 3D structures, represent a significant leap towards accurate mimicking of the lung’s cellular architecture, facilitating detailed drug and toxicity testing (5,6). However, the culture conditions are sophisticated and often require a dismantling of the natural architecture for infection. Finally, the lung-on-chip model incorporates a multicellular tissue environment with electronic monitoring systems for studies of complex interactions under flow conditions but the set up and validation remains challenging (7,8).

Precision-cut lung slices (PCLS) are an easy and physiological tool in respiratory research, providing a 3D *ex vivo* platform that bridges the gap between *in vitro* models and *in vivo* studies. Since the late 1970s (9–11). PCLS have been used in diverse areas, such as toxicology (12,13), drug and vaccine evaluation (14,15), lung physiology (16–18), and studies of lung diseases, including obstructive (19,20) and fibrotic conditions (21). All resident cell types and their intercellular interactions are maintained in these slices (22,23), in which the native three-dimensional architecture and complex microarchitecture of the lung, encompassing both airway and vascular components, are preserved (24,25). This preservation represents a marked improvement over simpler systems, such as submerged models and air-liquid interface cultures, which do not fully capture the intricate tissue structure and cellular interactions of the lung. Unlike sophisticated systems, such as the lung-on-chip, which requires careful setting up and calibration, PCLS are relatively straightforward to prepare and are suitable for immediate use for multiple applications.

PCLS can, thus, be used+ for detailed studies of cellular responses to various stimuli, including inflammation and infection, in a controlled environment that mirrors the *in vivo* setting without the associated ethical concerns of animal experimentation. Recent advances in single-cell sequencing have revealed the presence of both innate and adaptive immune cells in PCLS (26), confirming the ability to simulate complex immune responses in this model. PCLS are amenable to proteomics, metabolomics and transcriptomics studies (27). Their viability is typically limited to about 7-14 days, posing challenges for long-term studies (18).

Refinements of the protocols for generating PCLS over the years have extended the viability and functionality of these tissue slices (28), paving the way for more extensive investigations into cellular functions and phenotypes. Optical microscopy techniques have complemented these efforts, enabling researchers to visualize cellular dynamics and structural changes in real-time, thereby providing deeper insight into respiratory physiology and pathology (29).

Precision-cut lung slices (PCLS) have emerged as a key tool in the study of infectious diseases, providing a unique *ex vivo* model for exploring the molecular mechanisms of infection and evaluating antimicrobial drugs. The PCLS technique has been developed in various animal species (30,31) for the study of diseases, such as bovine tuberculosis (32), influenza (30,33), or respiratory syncytial virus (RSV) infection (34,35). The use of PCLS makes it possible to reduce the number of animals used, in accordance with the 3Rs rule and ethical requirements to decrease the use of live animals (36). Human PCLS have also been used to investigate infections caused by intracellular bacteria, as *Chlamydia pneumoniae* (37), *Legionella pneumophila* (38), and *Coxiella burnetii* (39).

Further studies expanded the use of PCLS to include a broad range of human pathogens, particularly those for which traditional animal models do not mimic human disease accurately due to host-specificity issues (40). PCLS are also particularly valuable for studies of co-infections with two viruses (41), two bacteria (42), or for comparing viral and bacterial infections in parallel (34). PLCS also constitute a very useful model for studying pathogens in situations in which no suitable animal model is available or animal models are challenging and costly to use, particularly for pathogens like SARS-CoV-2 and *Mycobacterium tuberculosis* (Mtb), which can be manipulated only in biosafety level 3 conditions or above (32,43).

For SARS-CoV-2, the virus responsible for the COVID-19 pandemic, in particular, PCLS have proved an invaluable resource (44,45). This model has been used to predict susceptibility to SARS-CoV-2 in different animal species (46), minimizing the need for live animal testing and overcoming the problem of an absence of suitable animal models early in the pandemic. This approach has not only shed light on the pathogenesis of COVID-19, but has also facilitated direct comparisons of different variants of the virus, thereby enhancing our ability to assess the efficacy of various antiviral compounds. For the human pathogen Mtb, murine PCLS have been used to study the changes in alveolar morphology and the induction of TNF-α; these studies validated this system as an alternative model for studies of different aspects of tuberculosis pathogenesis under controlled conditions mimicking as closely as possible the events *in vivo* (43). We previously compared the responses of bovine PCLS to Mtb and *Mycobacterium bovis* (32) but human PCLS have never before been used to study Mtb.

Various protocols for PCLS preparation have been published (32,47,48), but they do not provide sufficient details for the preparation of PCLS. Moreover, confocal microscopy have been widely used to observe PCLS (29,49) including live cell imaging. However, by overcoming several limitations of previously published research, our study provides significant advances in microscopy techniques. While previous studies provided only a brief overview of immunofluorescence techniques, our research offers a comprehensive and detailed step-by-step explanation, enhancing reproducibility and clarity in methodology for other research teams wishing to implement this protocol. Additionally, in contrast to previous studies that predominantly concentrated on the identification of cellular markers, our research extends this methodology to encompass the detection of pathogen markers within infected cells, a feature that is seldom described in the context of infectious agents within the PCLS model. Whereas other studies were constrained to the utilisation of non-human PCLS, the detection of these pathogens in the context of human PCLS paves the way for the utilization of this model to investigate major infectious agents in public health. Moreover, while previous studies have frequently presented images of PCLS in regions exhibiting collapsed alveolar architecture, our methodology emphasizes the preservation of connective tissue architecture with well-expanded alveolar lumina, thereby ensuring the closest possible resemblance to the structural characteristics of the in vivo alveolar microenvironment. Finally, we discuss the limitations of earlier studies that were restricted to the tissue and cellular levels without high-resolution imaging capabilities. The present study represents the first description of the utilization of transmission electron microscopy in the context of PCLS, whereby detailed visualization of subcellular architecture and organelles Thus, the protocol described here provides a detailed description of the infection and imaging steps, can be readily adapted to specific experimental requirements, and is easy to implement by users with experience in tissue culture.

This protocol facilitates the observation of intracellular pathogens and their impact on cell architecture by allowing a detailed analysis of the alveolar microenvironment infected with SARS-CoV-2 or Mtb, from tissue to subcellular levels, through confocal imaging and transmission electron microscopy (TEM). High resolution imaging of Mtb infected PCLS revealed that mycobacteria were not only found in alveolar macrophages, but also localized within the alveolar wall, embedded in the connective tissue and, on occasion, in close proximity to an alveolar epithelial cell. PCLS are an asset to study early events occurring in the alveoli, such intimate contacts and the role of lung epithelial cells in TB are of great interest but still overlooked (50). For SARS-CoV-2, the present study is the first to demonstrate, using confocal microscopy, that different variants of the SARS-CoV-2 virus can enter cells and initiate a viral life cycle in the alveoli. This finding is of considerable significance, as it provides further evidence that, despite the effects of genetic drift and spike mutations, the SARS-CoV-2 retains the capacity to directly target alveolar cells, in addition to the damage caused by the activation of innate immunity.

## Materials

### Biological materials

□ Surgical resection fragment of human lung

The pathologist excised this fragment from a surgical specimen of the lung, which may have been a lobectomy, a segmentectomy, or a wedge resection.

▲**Caution:** The surgical specimen must be transported extemporaneously from the operating room to the pathology laboratory. The surgical resection fragment must be kept at 4°C in non supplemented RPMI medium in a falcon tube until use and should be manipulated in an appropriate biosafety level cabinet and laboratory (BSL2 or BSL3 depending on whether the donor had an infectious disease).

▲**Critical:** We recommend using lung tissues within six hours of their excision during surgery. Sample quality has proved critical and we recommend the evaluation of emphysema and fibrosis by histology.

#### PCLS preparation

##### Reagents

- Dulbecco’s phosphate-buffered saline, 1X, pH 7.0, calcium-and magnesium-free, for cell culture (Sigma-Aldrich, cat. no. D8537-500ML)
- Decomplemented Fetal bovine serum (FBS) (Sigma, F7524)
- Roswell Park Memorial Institute medium (RPMI) (Gibco, cat. no. 31870025)
- L-glutamine or GlutaMax supplement (Gibco, cat. no. 35050038)
- Antibiotic cocktail: penicillin/streptomycin (Thermo Fisher Scientific, cat. no. 15140122) or polymyxin-B, amphotericin-B, nalidixic acid, trimethoprim, and azlocillin (PANTA^TM^), (BD BBL^TM^, cat. no. 245114)
- UltraPure^TM^ low-melting-point agarose (Invitrogen, cat. no. 16520050)

▲**Critical** We recommend using UltraPure^TM^ low-melting point agarose rather than other matrices to fill the lung biopsy specimen. We compared this reagent with gelatin, which is known to dissolve more efficiently at 37 °C, but we found that the alveolar sacs were less well expanded than with low-melting point agarose and that they appeared collapsed with gelatin (Figure S1, Figure 2e and 2f)

**Figure 1:**
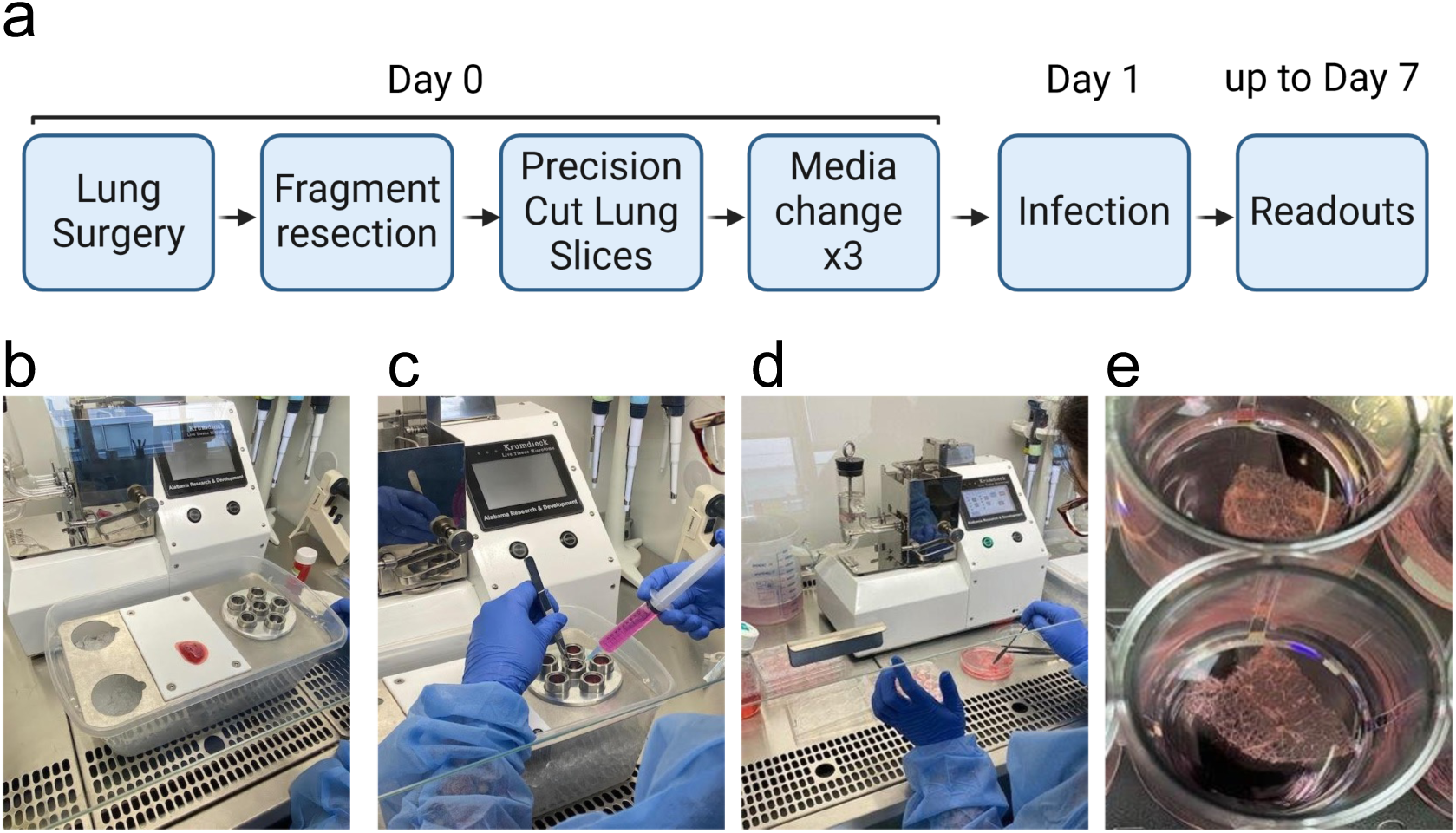
Schematic diagram of workflow and timeline. a) PCLS must be obtained from a fresh lung fragment, and the medium must be replaced at least three times before leaving the sample to rest overnight. We recommend infecting the PCLS the following morning, and we obtained convincing results for infections performed up to seven days after the PCLS was obtained. b) Lung fragment after resection, photographed on the working surface of the tissue-embedding unit. This fragment was cut into smaller pieces and placed in the plunger chambers. c) While injecting the low-melting point agarose with a syringe fitted with a needle, hold the lung piece in the plunger chamber with tweezers. d) Recover the PCLS in a Glass petri dish containing supplemented RPMI, select the best PCLS and place them in culture plates e) PCLS in a 24-well tissue culture plate. Representative images are shown from one experiment of 12 performed.

**Figure 2:**
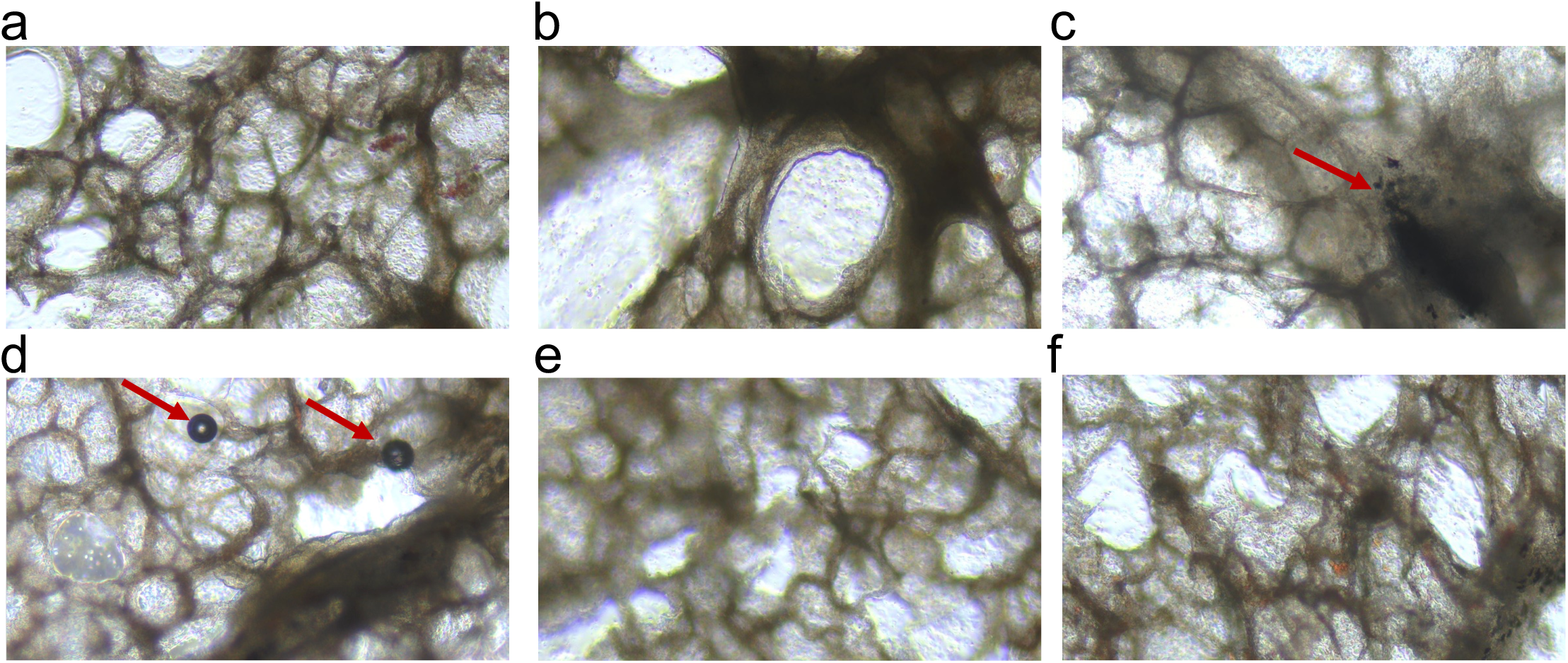
Light microscopy images of human PCLS. PCLS were obtained as described in figure 1 and observed under a light microscope (enlargement x40). (a) PCLS containing numerous alveoli and (b) a few bronchioles (thick and wavy epithelium). (c) Lung tobacco damage in a smoker. (d) Air bubbles trapped in low-melting point agarose can sometimes be visualized before the completion of the washing steps. (e, f) Collapsed alveolar structures observed when gelatin is used to fill the lung biopsy specimen. Representative images from one experiment of 12 performed are shown, except in e) and f) in which the results of one of two experiments are shown.

##### Equipment

- Alabama R&D tissue slicer (formerly known as the Krumdieck tissue slicer) (Alabama Research & Development, cat. no. MD6000) and at least one of its accessories: the tissue-embedding unit.

▲**Critical**: We recommend using the Alabama R&D tissue slicer to prepare aseptic, thin slices of living tissues. The biopsy specimen is embedded in a device with a plunger, which is moved by an arm, and a blade cuts the tissue. The speeds of both the arm and the blade can be modified. We tested another device (Leica Vibratome), which we found to be less satisfactory as cutting was slower, it was more difficult to cut outside of the parenchyma and the instrument could be cleaned and disinfected but not sterilized.

- Double-edged safety razor blades (such as Gillette Platinum)
- CO_2_ incubator, set at 37 °C and 5% CO_2_
- Biological safety cabinet
- Water bath, set at 39 °C
- Glass beaker, sterile, 100 mL volume (optimal)
- Microwave oven (700 to 900 W)
- Precision scale, resolution 0.001 g (optimal)
- Corning Co-star 24-well tissue culture (TC) plates (Thermo Fisher Scientific, cat. no. 353047)
- Inverted microscope for tissue culture
- Pipette controller, Pipet-Aid (such as Drummond^TM^, Thermo Fisher Scientific, cat. no. 11714819)
- 10 mL graduated serological pipettes, sterile (such as BD Falcon^TM^, cat. no. 357551)
- Gilson pipette set (P20, P200, P1000)
- Filter tips, sterile
- Dissection kit (scalpel, scissors, and tweezers)
- 2.5 mL syringe (BD Biosciences, cat. no. 334031)
- 23 G needles (BD Microlance 3)
- Sharpsafe ® container and biohazard bin
- Glass petri dishes, sterile

#### Infection of PCLS with SARS-CoV-2 (option A)

##### Reagents

- SARS-CoV-2 Wuhan strain (GISAID EPI-ISL-16833248), B.1.617 strain (Delta) and B.1.1.529 strain (Omicron BA.1), (tittered, in our study we used viral particles produced as described in Maquart et al., Photochemical & Photobiological Sciences, 2022 (51))
- Roswell Park Memorial Institute medium (RPMI) (Gibco, cat. no. 31870025)
- Decomplemented Fetal bovine serum (FBS) (Sigma, F7524)
- Trypsin (Gibco, cat.no. 11560626)
- Antibiotic cocktail: penicillin/streptomycin (Thermo Fisher Scientific, cat.no. 15140122)

##### Equipment

- CO_2_ incubator, set at 37 °C and 5% CO_2_
- Biological safety cabinet
- Gilson pipette set (P20, P200, P1000)
- Filter tips, sterile

#### Infection of PLCS with *Mycobacterium tuberculosis* (Mtb) (option B)

##### Reagents

- Mtb strain (tittered; in our study we used HN878-GFP strain, produced as described in Doz-Deblauwe et al., LSA 2024 (52))
- 7H9 broth (BD Difco^TM,^ ref 271310)
- Middlebrook ADC Enrichment (BD Dico^TM^ BBL^TM^, ref 211887)
- Tween 80, (Sigma, ref P4780)
- 7H11 agar base (BD BBL^TM,^ ref 212203)
- L-asparagine (Sigma, ref A4159)
- Middlebrook OADC Enrichment (BD Dico^TM^ BBL^TM^, ref 211886)
- Glycerol (Carlo Erba, ref 245114)

##### Equipment

- Sterile Glass petri dishes, 55 mm in diameter, with 3 vents (Grosseron GRS12104)
- Gilson pipette set (P20, P200, P1000)
- Filter tips, sterile
- CO_2_ incubator, set at 37 °C and 5% CO_2_
- Biological safety cabinet
- Centrifuge

#### PCLS imaging with a confocal microscope

##### Reagents

- Dulbecco’s phosphate-buffered saline, 1X, pH 7.0, calcium- and magnesium-free, for cell culture (Sigma-Aldrich, cat. no. D8537-500ML)
- Human Fc receptor binding inhibitor (eBioscience, ref. 14-9161-71)
- Horse serum (Sigma, H1270)
- Triton X100 (Sigma, T8787)
- Paraformaldehyde (such as Paraformaldehyde 32% solution, EM Grade, Electron Microscopy Science, cat. no.15714) ▲**Caution:** Irritant and carcinogenic. Read and understand safety data sheet before use.
- Primary and secondary antibodies (all the antibodies and dyes used in our study are listed in the table below)

| Reactivity | Isotype | Clone | Supplier, reference | Conjugation | Recommended concentration |
| --- | --- | --- | --- | --- | --- |
| Human Fc receptor binding inhibitor | Polyclonal | - | eBioscience, 14-9161-71 | - | 20 µg/mL |
| Anti-human CD163 | Mouse IgG1 | EDHu-1 | Biorad, MCA1853T | - | 5 µg/mL |
| Anti-mouse IgG1 | Goat IgG | - | Invitrogen, A21240 | AlexaFluor<br>647 | 10 µg/mL |
| Anti SARS-Cov-2<br>nucleoprotein | Mouse<br>IgG2ak | 1C7C7 | Sigma,<br>ZMS1075 | - | 1 µg/mL |
| Anti-mouse IgG | Goat | - | Thermo Fisher<br>Scientific,<br>#P-852 | PE | 10 µg/mL |
| Hoechst 33342<br>(counterstain) | - | - | Thermo Fisher<br>Scientific, 62249 | - | 10 µg/mL |

##### Equipment

- Confocal microscope (such as the Leica SP8 spectral confocal microscope)
- µ-Dish, 35 mm diameter, glass coverslip base #1.5H (81158, Ibidi)

#### PCLS imaging by TEM

##### Reagents

- Electron microscopy fixative mixture as described by McDowell and Trump (4% paraformaldehyde + 1% glutaraldehyde)

▲**Caution:** Irritant and carcinogenic. Read and understand safety data sheet before use.

- Sorensen’s phosphate buffer washing solution
- Epon embedding resin
- Lead citrate
- DPBS (Gibco, ref 18192-014)
- Paraformaldehyde (EM grade, ref GP26-250)
- Glutaraldehyde (VWR, ref 0875-100 mL)
- NaOH pellets (Carlo Erba, ref 480507)
- Sodium phosphate monobasic monohydrate (NaH_2_PO_4_·H_2_O) (Sigma, ref 1.06346.05.00)
- disodium phosphate (Na_2_HPO_4)_ (VWR, 280-26.292)
- NaCl (Merck, ref 1.06404.1000)
- Distilled water
- Osmium tetroxide (EMS, ref 19-190)
- LX112 resin (Sigma, ref 45-345-1L-F)
- Dodecenyl succinic anhydride (DDSA) (Sigma, ref 45346-250 mL-F)
- Propylene oxide (Thermofisher scientific, 03765-K2)
- Nadic methyl anhydride (NMA) (Sigma, ref 45347-1L-F)
- 2,4,6-trisdimethylaminomethylphenol (DMP30) (Sigma, ref T58203-50 mL)
- sodium borate (Formula: Na_2_B_4_O_7_·10 H_2_O) (Merck, ref 1.06308.05.00)
- toluidine blue (Sigma, ref T3260-256)
- uranyl acetate (formula: (CH_3_COO)2UO_2_·2H_2_O) (Merck, ref 8473)
- lead nitrate (formula: Pb(NO_3_)_2_) (Sigma, ref 1.07398.0100)
- trisodium citrate (formula: Na_3_C_6_H_5_O_7_·2H_2_O) (VWR, ref 27833.237)
- mineral oil (Cooper)
- Uranyl acetate (Merck, ref 8473)
- sterile injectable water (Fresenius, ref B230531)
- Ethanol (aqueous solutions at 50%, 70%, 90%, 100%) (VWR, ref 83813.360)

##### Equipment

- Ovens (Memmert)
- Laboratory fume hood (Sidpa 72, EN 14175)
- Flask (Duran, 1L)
- Light microscope (Nikon eclipse E200)
- Ultramicrotome (Leica, EMUC7)
- Block-trimming device (pyramitome) (Leica EMTRIM)
- Heated magnetic stirrers (IKA RH Basic 2)
- pH meter (Eutech instrument PH510)
- Precision balance (Sartorius, LP 1200S)
- 500 mL, 2000 mL Graduated flask (Duran)
- Graduated cylinders (Duran)
- Beaker (Duran)
- Pasteur pipette (Copan, 091-E14)
- 50 mL Falcon tube
- 20 mL syringes (BD, 300296)
- Magnetic stir bars (Merck)
- Weighing boats (Scientific Heathrow HS-1424B)
- Filter membranes with 0.20 µm pores (Millex, SLGS03355)
- Low-ash (max 0.06%) filter paper composed of cotton fibers (Prat Dumas, 150 mm)
- Quantitative filter paper (VWR, ref 516-0853)
- Lead pot
- Glass petri dish (Anumbra, 120 mm)
- Gelatin capsule (EMS, ref 70103)
- Capsule holder (Agar scientific, G3530)
- Perfect Loop for electron microscopy (Diatome)
- Electron microscopy MESH grids (Delta microscopy, ref G200-Au) and single-hole grids (EMS, ref G210-Ni)
- Electron microscopy grid box (Leica, 16705525)
- Aluminium foil
- Hot plate
- Parafilm (Sigma, ref P7793)
- Transmission electron microscope (such as JEOL 1400, Tokyo, Japan)

##### Reagent setup

**Low-melting point agarose** must be prepared immediately before use. Weigh the desired amount of low-melting point agarose powder with a precision scale. The final concentration in the medium should be 1.5%. Add the powder to a sterile glass beaker together with the correct volume of RPMI medium and heat in a microwave. We recommend heating for few seconds (20-30 s), then stopping, gently mixing the volume in the beaker and heating again. These steps should be repeated until the low-melting point agarose is completely dissolved. The beaker should then be transferred, with caution, to a water bath set at 39 °C.

▲**Caution:** The agarose may be very hot after heating in the microwave oven and the beaker should therefore be handled with heat-resistant gloves.

▲**Critical:** As the melting point of the agarose is low (37 °C), do not heat for too long and stay in front of the microwave over to monitor the melting process. Bubbles and foam may form rapidly.

**Supplemented RPMI medium** can be prepared several days in advance.

L-glutamine or Glutamax^TM^ should be added to a final concentration of 2 mM, together with 10% heat-inactivated decomplemented FBS.

**For PCLS generation steps :** a cocktail of antibiotics/antifungal agents can be added to the medium to prevent contamination. The choice of cocktail should reflect the final use of the PCLS. Penicillin/streptomycin mixtures are often used (final concentrations 100 IU/mL and 100 µg/mL, respectively) but should be replaced by PANTA cocktail (final concentration XXX µg/mL) for studies of mycobacterial infections.

**For PCLS infection steps :** do not add antibiotics to the RPMI medium.

▲**Caution:** The manufacturer’s instructions should be followed for reconstitution of the antibiotic mixture from the lyophilized powder in the vial.

▲**Critical:** Once reconstituted, antibiotic mixtures should be used within 72 h if stored at 2–8 °C, or within six months if stored at temperatures at or below -20 °C.

**Paraformaldehyde 4%** is obtained after 8 times dilution with DPBS, from the 32% solution.

▲**Caution:** Irritant and carcinogenic. Wear personal protective equipment (PPE) to prevent contact with the skin and eyes. Prepare and handle the solution in a laboratory fume hood to prevent inhalation. Paraformaldehyde waste should be placed in a tightly sealed, appropriately labeled hazardous waste container.

##### Electron microscopy fixative mixture as described by McDowell and Trump

Preparation of 40% paraformaldehyde: Add 200 mL distilled water and 80 g paraformaldehyde to a 500 mL flask and heat for 48 hours at 80 °C in an oven. While hot, add a few drops of 1 N NaOH to clear the solution, which should then be left to cool and filtered (ashless filter).

Preparation of the fixative mixture: The following should be added to a graduated flask, in the following order: NaH_2_PO_4_·H_2_O (23.20 g), NaOH (5.40 g), distilled water (1500 mL), 40% paraformaldehyde (200 mL) and 50% glutaraldehyde (40 mL). Adjust the volume to 2000 mL with distilled water. Adjust the pH to 7.35 with 1 N NaOH (use a pH meter for this). Store at +4 °C.

▲**Caution:** Irritant and carcinogenic. Wear personal protective equipment (PPE) to prevent contact with the skin and eyes. Prepare and handle the solution in a laboratory fume hood to prevent inhalation. Paraformaldehyde/glutaraldehyde waste should be placed in a tightly sealed, appropriately labeled hazardous waste container.

##### Sorensen’s phosphate buffer (0.3 M) and washing solution

Preparation of 0.3 M disodium phosphate (Na_2_HPO_4_, M= 141.96 g/mol) Add 6.39 g to 150 mL sterile injectable water and dissolve on a heated magnetic stirrer at 30 °C with a magnetic stir bar.

Preparation of 0.3 M monosodium phosphate (NaH_2_PO_4_·H_2_O M = 137.99 g /mol): Add 3.10 g to 75 mL sterile injectable water and dissolve on a heated magnetic stirrer at 30 °C with a magnetic stir bar.

Preparation of phosphate buffer: Stir the disodium phosphate solution on a magnetic stirrer and insert the pH meter. Slowly add monosodium phosphate solution until a pH of 7.3 is reached. Store for a maximum of 10 days at room temperature in a closed bottle.

Preparation of washing solution: In a glass bottle, mix 75 mL Sorensen’s phosphate buffer, 75 mL sterile water and 0.3 g NaCl.

▲**Caution:** Irritant and carcinogenic. Wear personal protective equipment (PPE) to prevent *any* contact with the skin and eyes. Prepare and use the solution in a laboratory fume hood to prevent inhalation. Paraformaldehyde/glutaraldehyde waste should be placed in a tightly sealed, appropriately labeled hazardous waste container.

##### Osmium tetroxide (2%)

Mix equal volumes of 0.3 M Sorensen’s phosphate buffer and 4% osmium tetroxide 4%.

▴**Caution:** Solid OsO_4_ is volatile and vaporizes readily from aqueous solution even at room temperature. High acute toxicity; severe irritant of the eyes and respiratory tract. The vapor can cause serious eye damage. May cause organ damage (CNS, eyes, skin, kidney). Prepare and handle solutions in a laboratory fume hood. Wear personal protective equipment (PPE) to prevent *any* contact with the skin and eyes. Store pure osmium tetroxide and its concentrated solutions in appropriate, sealed glass containers within unbreakable secondary containment. Osmium-containing waste should be placed in a tightly sealed, appropriately labeled hazardous waste container.

##### Epon embedding resin

Preparation of Epon medium A: Mix 62 mL LX112 resin and 100 mL dodecenyl succinic anhydride (DDSA). Mix, with stirring for 2 hours, and transfer to a 50 mL Falcon tube. Store at +4 °C.

Preparation of Epon medium B: Mix 100 mL LX112 resin with 89 mL of nadic methyl anhydride (NMA). Stir for 2 hours on a magnetic stirrer and then transfer to a 50 mL Falcon tube. Store at +4 °C.

Preparation of EPON embedding medium: Mix Media A and B as specified in the manufacturer’s instructions (table below) and stir for 1 hour on a magnetic stirrer. Add the necessary amount of 2,4,6-trisdimethylaminomethylphenol (DMP30) accelerator to achieve a proportion of 1.5%, as specified in the manufacturer’s instructions and stir for 15 minutes on a magnetic stirrer, before placing at 56 °C for 15 minutes to remove residual bubbles. Once the embedding medium has been used, any surplus can be stored in 20 mL syringes at -20 °C.

| Total volume<br>(mL) | Medium A<br>(mL) | Medium B<br>(mL) | DMP30<br>(mL) |
| --- | --- | --- | --- |
| 4.5 | 2.25 | 2.25 | 0.06 |
| 9 | 4.5 | 4.5 | 0.15 |
| 15 | 7.7 | 7.7 | 0.22 |
| 30 | 15 | 15 | 0.45 |
| 45 | 22.5 | 22.5 | 0.675 |
| 50 | 25 | 25 | 0.75 |
| 60 | 30 | 30 | 0.9 |
| 80 | 40 | 40 | 1,2 |
| 100 | 50 | 50 | 1.5 |

▴**Critical:** Must be prepared on the day of use. The medium should be brought to room temperature before use.

##### Toluidine blue

Mix 200 mL sterile water and 2 g sodium borate (Formula: Na_2_B_4_O_7_·10 H_2_O - molecular weight = 381.4 g/mol) with stirring on a magnetic stirrer to dissolve the sodium borate.

Add 1 g toluidine blue and allow it to dissolve. Filter through low-ash (max. 0.06%) filter paper composed of cotton fibers and store in 20 mL plastic syringes. Fit the syringe with a Millipore 0.20 µm-pore filter before use.

##### Uranyl acetate

Mix 1.5 g uranyl acetate (formula: (CH_3_COO)2UO_2_·2H_2_O; molecular weight: 424.15 g/mol) with 30 mL sterile water (to obtain a final concentration of 5%). Dissolve by stirring with a magnetic stirrer in the dark. Store at + 4 °C in a glass bottle housed in an opaque lead pot (shelf life: up to 6 months).

▲**Caution:** Highly toxic material with low-level radioactivity. Wear personal protective equipment (PPE) to prevent any contact with the skin and eyes. Uranyl acetate residues and contaminated materials must be processed as radioactive waste.

▴**Critical:** Keep uranyl acetate away from the light whether in its dry crystal form, or in the form of the stored aqueous solution and during its use for staining. Store in an opaque container and let the solution stand for one day before use.

##### Lead citrate

Preparation of lead nitrate (formula: Pb(NO_3_)_2_; molecular weight: 331.2 g/mol): For a 1 M solution, dissolve 33.12 g lead nitrate in 100 mL sterile water. Stir and then transfer to a glass bottle (shelf life: up to 1 year)

Preparation of trisodium citrate (formula: Na_3_C_6_H_5_O_7_·2H_2_O; molecular weight: 294.1 g/mol): For a 1 M solution, dissolve 29.41 g trisodium citrate in 100 mL sterile water. Stir and then transfer to a glass bottle (shelf life: up to 1 year)

Preparation of sodium hydroxide (formula: NaOH; molecular weight: 40 g/mol): For a 1 M solution, dissolve 4 g NaOH in 100 mL sterile water. Stir and then transfer to a glass bottle (shelf life: up to 10 days)

Preparation of lead citrate

Mix 3 mL trisodium citrate and 16 mL sterile water in a 50 mL Falcon tube with a magnetic stirrer. Add 2 mL lead nitrate. Stir to obtain a milky solution. Add 4 mL sodium hydroxide (freshly prepared). Stir until a clear solution is obtained. Overlay the solution with mineral oil to prevent oxidation of the lead citrate. Store at room temperature (shelf life: up to 10 days)

▲**Caution:** Carcinogenic. Wear personal protective equipment (PPE) to prevent *any* contact with the skin and eyes.

## Procedures

### Phase 1: generation of PCLS

- **Time required: 1/2 day**

1. Dissect the lung biopsy specimen on the ice-cooled working surface of the tissue embedding unit. Cut it into 1 cm pieces (Figure 1).

▲**Caution:** Wear gloves and be careful not to cut yourself with the dissection instruments. Use a biohazard bin and Sharpsafe ® container to ensure the safe disposal of waste.

▲**Critical:** work at 4 °C. Carefully remove all the pulmonary pleurae with a scalpel blade. Incomplete removal can cause problems that will need to be resolved (see troubleshooting, below).

2. Use a syringe to transfer liquid RPMI supplemented with 1.5% low-melting point agarose to the mold plunger assembly blocks. Allow the agarose to solidify for 30 s before moving on to the next step.

▲**Critical:** The liquid must entirely cover the bottom of the plunger.

3. Place the pieces of lung into the mold plunger assembly blocks (1 per block).
4. Use a syringe fitted with a 23 G needle to fill each piece of lung with liquid RPMI supplemented with 1.5% low-melting point agarose. The lung pieces will swell and liquid will leak into the block. Continue to inject the medium until the block is full and then wait for the medium to solidify before moving on to the next step.

▲**Caution:** Manipulate the needle with caution and use a Sharpsafe ® container to ensure its safe disposal after use.

▲**Critical:** You may need to replace the needle if you fill have several blocks to fill, as the agarose solidifies rapidly and may block the needle. Use a tweezer to hold and orientate the piece of lung, which should at least double in size when the block is full.

◆ Troubleshooting

5. Set up the Alabama R&D Tissue Slicer equipment in accordance with the manufacturer’s instructions. Fill the reservoir with cold DPBS.

▲**Caution:** Manipulate the blade with caution and use a Sharpsafe ® container to dispose of it safely after use.

▲**Critical:** The temperature of the DPBS is crucial. If it is not sufficiently cold, the quality of the PCLS may be affected. We recommend changing the DPBS every 30 minutes, or more rapidly if room temperature is high.

6. Insert one plunger block containing a piece of lung into the Alabama R&D Tissue Slicer. Set the arm speed to 3/5 and the blade speed to 3/5 (default parameters) and begin cutting. ◆ Troubleshooting

▴**Critical:** The Alabama Tissue Slicer can be set to obtain PCLS of 100 to 1000 µm thick. For our study, we set the thickness to 100-150 µm.

7. Recover the PCLS in the glass trap.
8. Disperse the PCLS in a Glass petri dish containing supplemented RPMI medium (supplemented with L-glutamine, 10% decomplemented FBS, antibiotics).
9. Select the best PCLS and place them in 12- or 24-well TC plates (use a spatula or tweezers), with 1 PCLS and 1 mL supplemented RPMI medium per well.

▲**Critical:**If you use tweezers, make sure that they have blunt tips as tweezers with high-precision sharp tips may damage the PCLS.

10. Incubate the PCLS at 37 °C under an atmosphere containing 5% CO_2_.
11. Change the medium after 30 minutes, returning the PCLS to the incubator.
12. Repeat steps 10 and 11 (at least twice) to wash the PCLS and to remove the low-melting point agarose.
13. Observe the PCLS under a light microscope to ensure that the agarose has been removed
14. Incubate overnight at 37 °C under an atmosphere containing 5% CO_2_.

▴**Critical:**We recommend letting the tissue rest overnight to eliminate stress signals and to reverse any bronchoconstriction that may have occurred.

### Phase 2: Infection of the PCLS

- **Time required: 1/2 day**

15. Change the medium and observe the PCLS under a light microscope (Figure 2). If bronchioles are present in the PCLS, the ciliary activity of epithelial cells should be evident. Note that alveolar epithelial cells (pneumocytes) are not ciliated and it is therefore normal to observe no such movements in alveoli. ◆ Troubleshooting
16. Prepare your inoculum. Explanations for the two pathogens used in our study are provided below.

Option A: Infection with SARS-CoV-2

▲**Caution:** The handling of SARS-CoV-2 requires specific biosafety procedures and a biosafety level 3 laboratory.

i) Take a frozen vial of virus stock of known titer. Defrost the aliquot and pipette out the desired volume of the viral suspension for the addition of 1.3 x 10^6^ viruses per PCLS.
ii) Incubate the PCLS with the inoculum in RPMI medium supplemented with 1 µg/mL trypsin for 6 hours at 37 °C.
iii) Replace the medium with RPMI

medium supplemented with 10% decomplemented FBS and 1% antibiotic mixture (penicillin/streptomycin) and incubate at 37 °C overnight.

▲**Critical:** The use of trypsin significantly increases the efficiency of cell infection (through proteolytic cleavage of the spike envelope protein, increasing its affinity for the ACE2 entry receptor). In the absence of trypsin, the infection of PCLS by SARS-CoV-2 appears to be severely limited.

Option B: Infection with *Mycobacterium tuberculosis* (Mtb)

▲**Caution:** The handling of Mtb requires specific biosafety procedures and a biosafety level 3 laboratory.

i) Take a frozen vial of the mycobacterium of known titer. Defrost the aliquot and remove the desired volume of bacteria to achieve a density of 10^5^ CFU per PCLS.
ii) Incubate for 4-5 hours at 37 °C in 1 mL 7H9 medium supplemented with 10% ADC and 0.05% Tween 80 to allow the mycobacteria to reactivate.
iii) Centrifuge for 10 min at 1700 x *g* and room temperature.
iv) Carefully discard the supernatant and resuspend the pellet in the appropriate volume of supplemented antibiotic-free RPMI obtain 10^6^ CFU per mL.
v) Add 100 µL/10^5^ CFU per well.

▲**Critical:** We estimate that, 1 hour after infection, at least 10^4^ CFU have reached the PCLS and that this is sufficient to trigger PCLS responses (Remot et al., 2021 (32) and Figure S2). The size of the wells can be adjusted (use of 24- or 48-well TC plates) according to the size of the PCLS, to optimize infection.

vi) Check the inoculum by serial dilution on 7H11 agar plates supplemented with 10% OADC and 0.5 % glycerol. Incubate the plates for 2-3 weeks at 37 °C and then count the CFU.

### Phase 3: Imaging of the PCLS (confocal microscopy)

- **Time required: 1 day**

17. After infection, the PCLS should be fixed by incubation in 4% paraformaldehyde at 4 °C and then stored in DPBS at 4 °C. The incubation time for fixation must be calculated so as to ensure that the pathogens are inactivated. The fixation time determined for this protocol was overnight.

▲**Caution:** Paraformaldehyde is a carcinogenic, mutagenic and reprotoxic substance that must be handled in accordance with good health and safety practices. Read and understand the manufacturer’s safety data sheet carefully before use.

▲**Critical:** Fixed PCLS can be stored in DPBS at 4 °C for several months before imaging, provided that DPBS evaporation is prevented.

18. Place the PCLS with a spatula in a new plate at room temperature, choosing the plate format most appropriate for the size of your PCLS. Smaller wells are preferable, as they make it possible to optimize the amounts of antibodies used.

▲**Critical:** The volumes given below are those for a 48-well plate. They should be doubled for a 24-well TC plate.

19. Permeabilize the PCLS with 200 µL of 0.3% Triton X-100 in DPBS (30 min - 1 hour, at room temperature with gentle shaking) ◆ Troubleshooting

▴**Critical:** Vortex the DPBS-Triton solution before use.

20. Incubate the PCLS with 200 µL of DPBS supplemented with 0.1% Triton X-100, 10% horse serum and Fc block for 1 hour, at room temperature, with gentle shaking, to saturate the Fc receptors and prevent non-specific antibody binding.
21. Incubate the PCLS for at least 2 hours with 200 µL primary antibodies in DPBS supplemented with 0.1% Triton X-100 and 10% horse serum (at room temperature, with gentle shaking). ◆ Troubleshooting

PAUSE: you can choose to incubate overnight at 4 °C and then continue with the steps described below the following day.

22. Wash three times by replacing the medium with 500 µL DPBS and incubating for 5-10 min at room temperature.
23. Incubate the PCLS for at least 2 hours with 200 µL secondary antibodies diluted in DPBS supplemented with 0.1% Triton X-100 and 10% horse serum (at room temperature, with gentle shaking). ◆ Troubleshooting

PAUSE: you can choose to incubate overnight at 4 °C and then continue with the steps below the following day.

24. Wash three times by replacing the medium with 500 µL DPBS and incubating for 5-10 min at room temperature.
25. Incubate with Hoechst stain in DPBS for at least 30 minutes at room temperature..
◆ Troubleshooting
26. Wash for 5 min with 500 µL DPBS.
27. Place the PCLS in a 35 mm µ-Dish with a glass coverslip base for confocal imaging.

### Phase 4: Imaging of PCLS (TEM)

● **Time required: 2 days** (steps 28 to 42 : fixation to impregnation)
● **Time required: 2 days** (steps 43 to 45 : Embedding)
● **Time required: 1 day** (steps 46 to 69: Ultra-thin sections to staining with uranyl acetate and lead citrate)
28. After infection, place the PCLS in a microtube and add electron microscopy fixative (4% paraformaldehyde + 1% glutaraldehyde), ensuring that the sample remains immersed in the fixative. The duration of fixation must be calculated to ensure that the pathogens are inactivated (overnight at +4°C in this protocol).

▲**Caution:** These operations should be performed in a laboratory fume hood and with personal protective equipment (PPE). Waste paraformaldehyde/glutaraldehyde should be placed in a tightly sealed, appropriately labeled hazardous waste container.

▲**Critical:** samples must be fixed for at least 2 hours before the washing step.

Fixed PCLS must be kept at +4 °C and never at -20 °C to prevent the ultrastructural damages linked to freezing and thawing.

29. Wash the sample in Sorensen’s phosphate buffer washing solution: Empty the tubes by gradually tilting them, taking care to leave the sample at the bottom of the tube. Fill the tubes three-quarters full with the washing solution and then close them.
30. Change the washing solution every 45 minutes.
31. Repeat steps 28 and 29 three times (=4 washes in total)
32. After the last wash, store the tubes at +4 °C for a minimum of 12 hours.
33. Replace the washing solution with OsO_4_ solution (2% aqueous solution) as described above (1 mL per microtube) and place the tubes at room temperature in the dark for 1 hour.

▲**Caution:** These operations should be performed in a laboratory fume hood and with personal protective equipment (PPE). Osmium-containing waste should be placed in a tightly sealed, appropriately labeled hazardous waste container.

34. Rinse three times with sterile water.
35. **Dehydration** should be performed at room temperature: the samples should be dehydrated in ethanol baths of increasing concentrations and then in two baths of propylene oxide (see the table below).

36. **Pre-impregnation**: Thaw the syringes containing Epon resin at +4 °C, using 1 mL Epon per tube and 0.5 mL Epon per microtube.
37. In a Falcon tube, mix equal volumes (v/v) of propylene oxide and EPON resin from the syringe.
38. Replace the propylene oxide (from the final dehydration step) with the propylene oxide-Epon mixture (v/v).
39. Cover with aluminum foil and seal the tube.
40. Incubate the samples in the propylene oxide-Epon mixture for at least 2 hours at room temperature.
41. **Impregnation**: Thaw the syringes containing Epon resin at +4 °C, using 2 mL Epon per tube and 1 mL Epon per microtube.
42. Replace the mixture with the Epon resin from the syringe. Cover with aluminum foil without sealing the tube. Allow the samples to rest overnight at 4 °C in the Epon embedding medium to complete the impregnation.
43. **Embedding** is performed with gelatin capsules: Place the capsules in capsule holders, add 1 drop of EPON embedding medium, then the sample and fill the capsule to three-quarters full with EPON embedding medium.
44. Place the capsules in an oven at 60 °C for at least 48 hours to polymerize.
45. Before sectioning, trim the polymerized blocks into a pyramidal shape with a trapezoidal section, using the EM TRIM pyramitome to remove excess resin around the sample.
46. Cut **semi-thin sections** on an ultramicrotome with a histo-type diamond knife equipped with a water-filled trough for the collection of semi-thin sections.
47. Collect sections in a drop of water using collection rings and place them on a glass slide.
48. Dry the slide on a hotplate (60 °C).
49. Cover the sections with filtered toluidine blue.
50. Allow the sections to stain for 20 seconds without letting them dry out.
51. Rinse the sections with distilled water. Add a drop of ethanol if the stain is too concentrated and rinse.
52. Let the sections dry on a hotplate (60 °C).
53. Observe under a light microscope (Figure S3).
54. Select the area of interest for ultrathin sectioning. ◆ Troubleshooting
55. **Ultra-thin sections** are cut with ultra-type diamond knives equipped with a trough filled with water for collecting ultrathin sections.

|  | Number of baths | Time (minutes) |
| --- | --- | --- |
| 50% ethanol | 2 | 5 |
| 70% ethanol | 2 | 10 |
| 90% ethanol | 2 | 15 |
| 100% ethanol | 3 | 20 |
| Propylene oxide | 2 | 30 |

▲**Critical:** The following factors affect section quality: the angle of the knife relative to the block should be 6°, the cutting speed should be 1 mm/s, the water level in the knife trough must be perfectly adjusted (the shiny gray reflection of the knife should be visible), the large base of the section should be parallel to the knife edge. The ultrathin sections should be between 50 and 120 nm thick.

56. Collect sections with a Perfect Loop and place them on 3 mm diameter MESH grids made of gold, copper, or nickel.

▴**Critical:** It is recommended to use both MESH grids and single-hole grids to increase the chances of visualizing the areas of interest (alveolar tissue is a very loose tissue and the alveolar walls on ultrathin sections may lie below a bar of the grid and therefore not be visible.)

57. Place the grid on quantitative filter paper (with the sections facing up) in a Glass petri dish and allow it to dry at room temperature.
58. Store the grids in numbered specific grid boxes and note their location. The grid boxes should be opened and closed with caution.
59. **Staining with uranyl acetate:** Use a 1 mL syringe to pass the aqueous uranyl acetate through a Millipore filter with 0.20 µm pores into a plastic microtube.
60. Dilute 1/2 in absolute ethanol to contrast MESH grids. Use pure uranyl acetate for single-hole grids, then stir and place immediately in the dark.
61. Place the grids with the section side in contact with a pure uranyl acetate drop deposited on Parafilm in a Glass petri dish in the dark.
62. Leave in the dark for 5 to 10 minutes, depending on the contrast desired.
63. Rinse each grid in 2 drops of sterile water.
64. Dry the grids, section-side upward, on quantitative filter paper.
65. **Lead citrate-uranyl acetate staining:** Clean the plastic platform with 1 N NaOH. Place drops of lead citrate in a Glass petri dish on the plastic platform (in a chamber “saturated” with NaOH pellets).
66. Place the grids with the section side in contact with lead citrate drop for 5 to 10 minutes, depending on the desired contrast.
67. Rinse in dilute NaOH solution (1 mL of 1 N NaOH in 50 mL sterile water).
68. Dry the grids; section-side upward, on filter paper.
69. Observe the grids with a transmission electron microscope. ◆ Troubleshooting

**Troubleshooting:**

| Step | Problem | Possible reason | Solution |
| --- | --- | --- | --- |
| 4 | The lung section is not filled or only partially filled | The agarose was not injected properly | Check the agarose temperature, and inject again in a new plunger chamber |
| 6 | No PCLS obtained | The blade did not cut the lung section properly (the persistence of pleural tissues may be the source of the problem) | Make sure that the section is still in the plunger chamber, remove 1 mm of tissue with a scalpel and try again. Otherwise, prepare another piece of tissue.<br>You can also try increasing the slicer blade speed and decreasing the arm speed. |
| 14 | No ciliary activity observed | Agarose still present within the PCLS | Repeat washing steps |
| 18 | Tissue not permeabilized | • The Triton is not well dissolved in DPBS<br>• The incubation time was not long enough | • Vortex the solution before use<br>• Increase the incubation time, especially if the PCLS are thick |
| 20, 21 or 24 | No labeling observed | • The antibody is not binding<br>• The incubation time is not long enough | • Change antibody/validate your antibody<br>• Increase the incubation time (up to overnight at 4 °C) |
| 53 | Very few alveolar walls are visible on semi-thin sections relative to the numbers of bronchiolar structures, vascular structures and the amount of surrounding connective tissue | The initial part of the sample embedded in the resin block does not encompass the alveolar compartment, instead enclosing more proximal structures. | Continue to advance through the resin block by cutting new sections, to find the alveolar compartment and areas of interest |
| 68 | PCLS ultrastructure is not interpretable: all cells are lysed and the alveolar wall is disrupted | •PCLS fixation for TEM was insufficient<br>•PCLS were frozen and thawed | • Increase the duration of incubation in the fixative mixture (2 hours minimum)<br>• Fixed PCLS should be kept at +4 °C and never at -20 °C |
| 68 | Alveolar cells are barely distinguishable, with low contrast | Ultrathin section staining insufficient | Repeat the lead citrate and uranyl acetate staining step |
| 68 | Large electron-dense deposits are visible on the ultrathin section | Precipitates are present in uranyl acetate solution | Repeat the filtering of the uranyl acetate and then repeat the staining step |
| 68 | Alveolar walls are numerous on semi-thin sections (light microscopy) but barely visible on ultra-thin sections (TEM) | The bars of the MESH grids obstruct observations | Prepare single-hole grids in addition to MESH grids |
| 68 | Ultrastructural modifications are visible in infected conditions but difficult to link to the pathogen | Some ultrastructural modifications may be artifactual and limited to a single PCLS | Search for these ultrastructural modifications in a control uninfected PCLS processed in the same batch |

### Anticipated results

Our study cohort comprised 12 patients (see the table below), with a median age of 72 years (range: 57 to 84 years). The sex distribution was balanced, with equal numbers of female and male patients. The cohort included patients with a spectrum of pulmonary diseases, primarily adenocarcinomas with various histological patterns: acinar, papillary, and mixed. Two patients presented with metastatic disease due to extrapulmonary primary tumors, one ovarian and the other a colonic adenocarcinoma. The remaining diseases included one case of mixed pulmonary carcinoma with predominant squamous and minor neuroendocrine differentiation, and one case of granulomatous lesions. Surgical treatments differed between patients according to the characteristics of the tumor and included wedge resections, segmentectomies, and lobectomies targeting specific segments and lobes, respectively, for optimal disease management.

| Patient characteristics (n=12) |  |  |
| --- | --- | --- |
| Age | 71.5 ± 7.6 years (mean ± SD) | 57 - 84 years (range) |
| Sex | 6 women - 6 men | Sex ratio = 1 |
| Smoker | Non-smokers | 3 |
|  | Former smokers | 7 |
|  | Current smokers | 2 |
| Disease | Lung adenocarcinoma | 9 |
|  | Lung metastasis | 2 |
|  | Other | 1 |
| Surgical procedure | Lobectomy | 7 |
|  | Segmentectomy | 3 |
|  | Wedge | 2 |

Non-fixed lung resections were transported from the thoracic surgery unit to the pathology laboratory. Upon arrival, peripheral lung parenchyma samples were taken and placed in culture medium at 4 °C. The lung parenchyma was then fixed in 10% neutral buffered formalin by endobronchial insufflation and the instillation of formalin through a catheter. Surgical specimens were fixed for 24 to 48 hours, then subjected to overnight dehydration and embedding in paraffin for the preparation of tissue blocks. Sections 3 to 4 µm thick were cut from the blocks and stained with hematoxylin, phloxin and saffron. During pathology analyses for diagnosis, a centimeter-sized tissue fragment was excised from the peripheral non-tumor area. This fragment was subjected to histological analysis to assess fibrosis, inflammation and epithelial degeneration. Lung parenchyma typically had a normal architecture, with some subpleural dystrophic emphysematous lesions. The alveoli contained macrophages and bronchovascular axes were normal (Figure S4).

PCLS were prepared as described in the protocol above. We visualized extracellular and intracellular Mtb 48 hours post infection. Bacilli were taken up by CD163^+^ macrophages (Figure 3). Interestingly, despite *ex vivo* infection, Mtb was able to reach its target cells. Mtb within macrophages accounted for at least 25% of the observed Mtb events. Increasing the dose of Mtb did not trigger more phagocytic events, but Mtb appeared more clustered (data not shown). Mtb was able to replicate in PCLS, as demonstrated by the increase in CFU counts (Figure S2) and confocal images revealing the presence of several bacilli within each cell (Figure 3).

**Figure 3:**
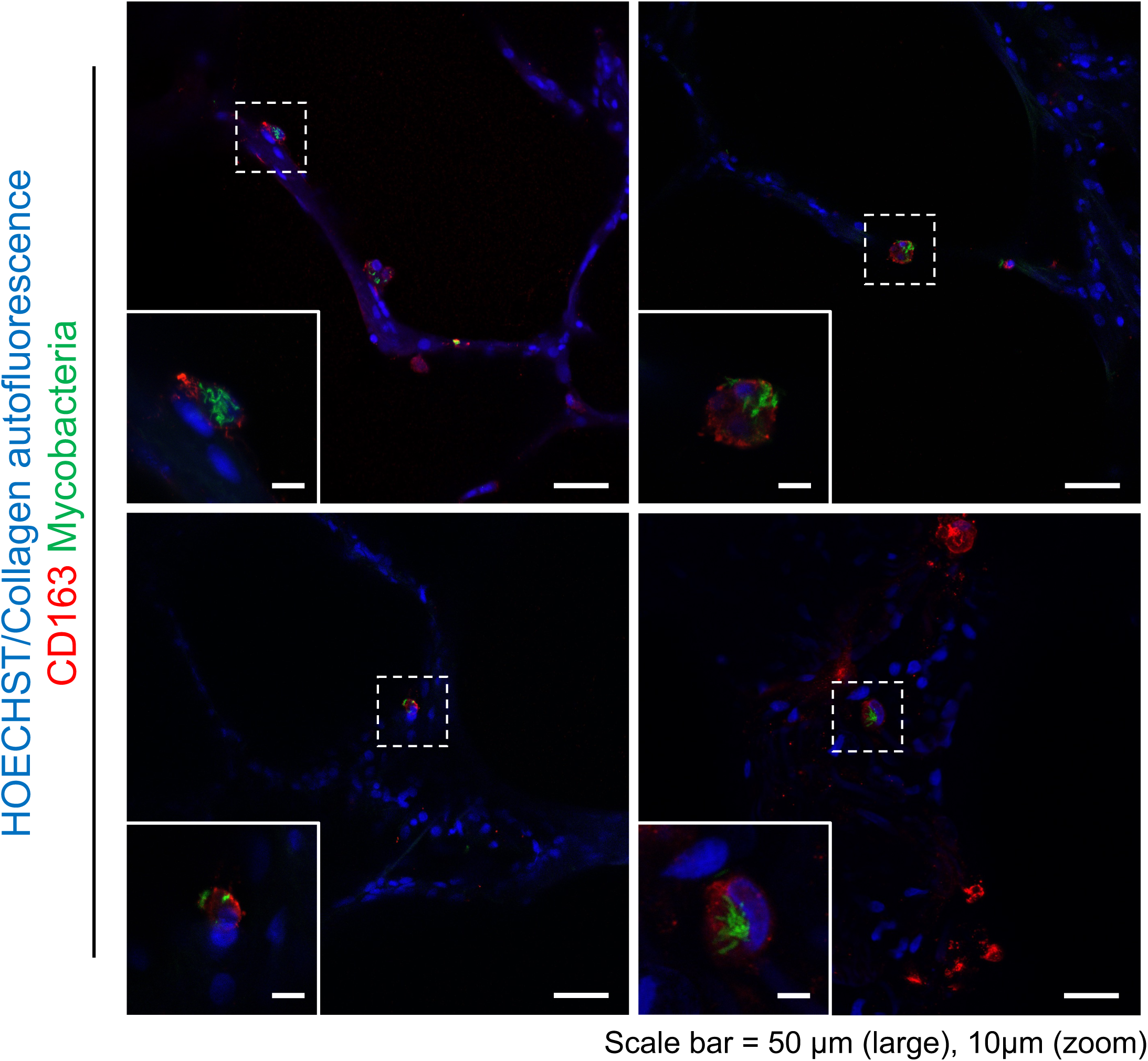
Localization *in situ* of *Mycobacterium tuberculosis* after the *ex vivo* infection of human PCLS. PCLS were obtained as described in figure 1. PCLS were infected with Mtb strain HN878-GFP and fixed with paraformaldehyde 2- or 5-days post infection. Fixed PCLS were stained with an anti-CD163 Ab (macrophages, detected with an AlexaFluor647-conjugated Ab, shown in red) and Hoechst stain (nuclei, blue). Representative images from one experiment of six performed are shown (1 patient per experiment, i.e., tested on 6 different patients).

It is notable that our study confirm that SARS-CoV-2 has the capacity to replicate in a tissular *ex vivo* model. A significant finding is that we observed cells infected with the Wuhan, Delta, and Omicron BA.1 strains of SARS-CoV-2. These infected cells had a morphology and distribution (with respect to the alveolar wall) consistent with alveolar epithelial cells and alveolar macrophages (Figure 4). Flattened elongated cells juxtaposed on the alveoli were observed, consistent with alveolar epithelial type I (AET1) cells, together with round protruding cells juxtaposed against the alveolar wall and consistent with alveolar epithelial type II (AET2) cells. Round cells free in the alveolar lumen and consistent with alveolar macrophages were also observed. Some PCLS (inter-patient variability) displayed a strong auto-fluorescence from connective fibers, with high levels of emission in blue and green wavelengths. We eliminated auto-fluorescence signals by acquiring confocal microscopy images in spectral mode. Fluorescence emission spectra were obtained for autofluorescence and the different fluorochromes used to facilitate the disentanglement of signals post-acquisition signals and the isolation of autofluorescence from signals of interest (Figure S5).

**Figure 4:**
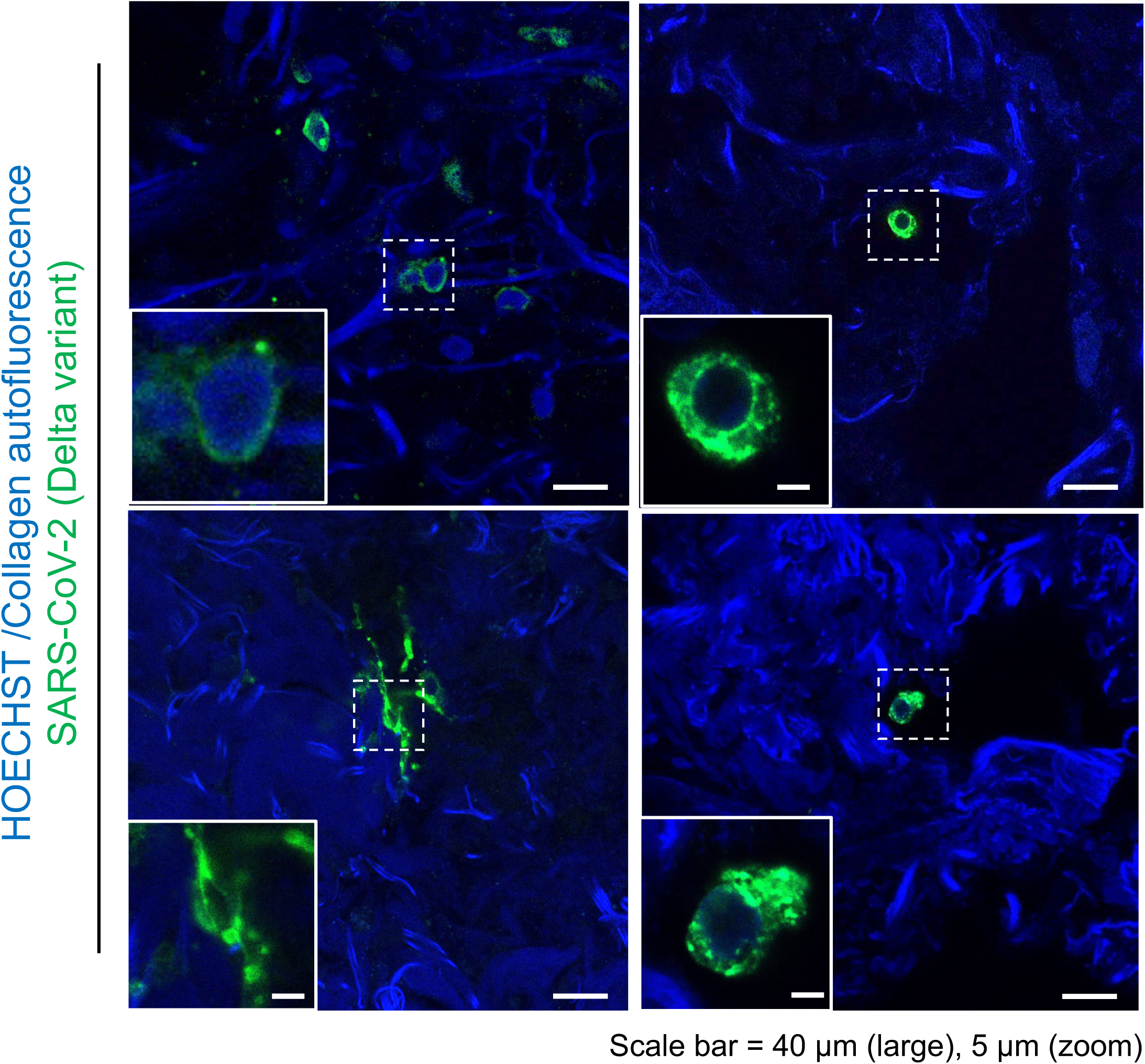
Localization *in situ* of SARS-CoV-2 after the *ex vivo* infection of human PCLS. PCLS were obtained as described in figure 1. PCLS were infected with the Delta strain and fixed with paraformaldehyde, 2 days post infection. Fixed PCLS were stained with anti-nucleocapsid Ab (detected with a phycoerythrin-conjugated Ab, shown in green) and Hoechst stain (nuclei, blue). Representative images from one of three experiments performed (1 patient per experiment, i.e., tested on 3 different patients).

Transmission electron microscopy (TEM) studies on uninfected PCLS demonstrated that ultrastructure was preserved in various cell types in the lung, up to one week after PCLS preparation (Figure 5, S6 and S7). The apical surfaces of ciliated cells carry microvilli and ciliary axonemes (Figure 5a). AET1 cells display their classic, flattened structure (Figure 5b), whereas AET2 cells appear cuboidal or spherical, with their cytoplasm densely packed with multilamellar bodies (Figure 5c). Alveolar macrophages can be distinguished on the basis of their indented nuclei and abundant cytoplasmic lysosomes and phagolysosomes (Figure 5d). The entirety of the blood-air barrier is also discernible, showing the epithelial basal lamina, capillary endothelial cells, and interwoven collagen and elastic fibers, all flanked by the slender cytoplasmic extensions of AET1 cells (Figure S6). Lung-resident immune cells, such as alveolar macrophages, granulocytes and lymphocytes, were also observed (Figure S7)

**Figure 5:**
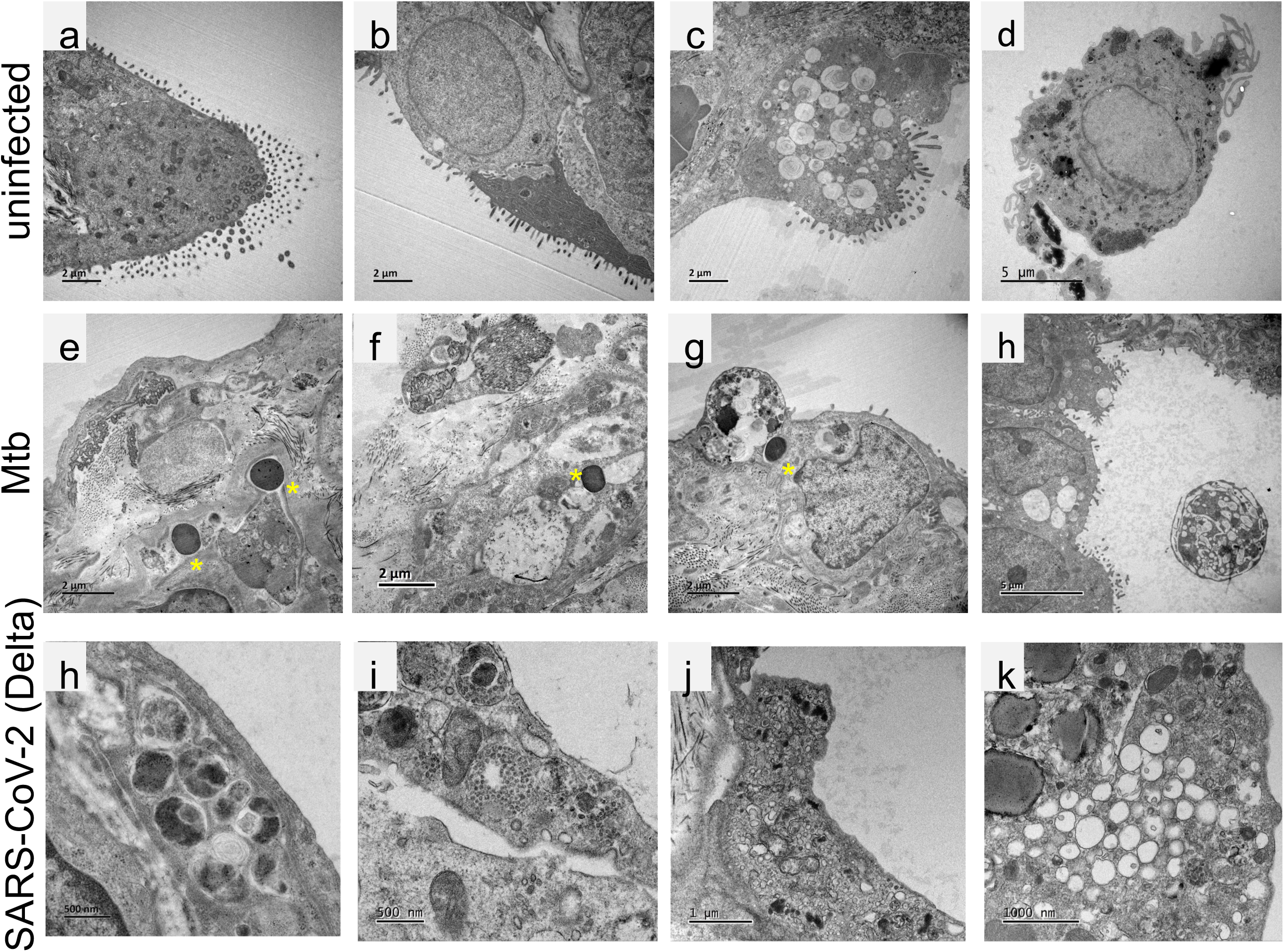
TEM. PCLS were obtained as described in figure 1. PCLS were fixed in 1% glutaraldehyde, 4% paraformaldehyde and embedded in Epon resin to obtain a block. Ultra-thin sections were cut from these blocks with an ultramicrotome, stained with 2% uranyl acetate and 5% lead citrate, and then observed with a transmission electron microscope (JEOL 1400). Representative images from one experiment of three performed are shown (1 patient per experiment, i.e., tested on 3 different patients).

In Mtb-infected PCLS, mycobacteria are readily identifiable as round (transverse section of the bacterium) to elliptical (longitudinal section of the bacterium) cells with an electron-dense cytoplasm (Figure 5e f and g). In some cases, an electron-lucent halo consistent with the glycosylated capsule was observed, clearly delineating the cell. The mycobacteria were found within the alveolar wall, embedded in the connective tissue and, on occasion, in close proximity to an alveolar epithelial cell. Ultrastructure was well preserved overall, but we observed apoptotic epithelial cells (Figure 5g, apoptotic AET2 cell above Mtb) and activated alveolar macrophages (Figure 5h).

TEM studies of SARS-CoV-2-infected PCLS also revealed a preservation of the ultrastructure of various cell types in the lung. Nevertheless, in infected PCLS, the ultrastructure is slightly altered, as evidenced by the detachment of some epithelial cells from the alveolar septum. From day 2 to day 5, we observed no assembly of complete viral particles in the cells. However, in alveolar epithelial cells, cell ultrastructure was significantly modified in infected PCLS relative to non-infected PCLS (Figure 5): multilamellar structures with electron-dense content consistent with autophagy-lysosomal elements (h), large round vesicles filled with many smaller round elements consistent with multivesicular bodies (MVBs) (i), a major increase in the number of secretory vesicles (j), and clusters of single-membrane vesicles (k). The observed changes may be induced by a viral infection of the cell, in this by SARS-CoV-2, but this cannot be concluded definitively without further investigations. PCLS brought a real step forward to study the initial steps of infection of host lung cells by two major respiratory pathogens that will help in the development of new COVID-19 or TB treatment strategies. This model is easily adaptable to respond to new and emerging outbreaks.

**Regulatory Aspects:** This research project is a collaborative effort between the University Hospital Center of Tours (CHU) and the French National Research Institute for Agriculture, Food, and Environment (INRAE, Unit UMR1282 Infectiology and Public Health), in association with the French National Institute of Health and Medical Research Unit U1259 MAVIVH. The partnership is formalized under the C96-02 agreement. The research project have been approved by the ethics committee of Tours University Hospital. The project involves the use of a biological collection related to pulmonary cancers, referenced under declaration number DC-2008-308. This collection has been duly declared in compliance with the regulatory requirements governing the use of human biological samples for research purposes. The collection of data from patients has been declared to the French Data Protection Authority (*Commission nationale de l’informatique et des libertés*).

## Author contributions

AR and SE set up the protocols, performed experiments, analyzed the data, obtained funding for this project, prepared figures and wrote the manuscript. SE managed reglementary statements, ethical supervision of the study, patient inclusion and patient data collection. AB and MM performed SARS-CoV-2 experiments; FCa, MSV, EDD and BB performed Mtb experiments. FCa generated and prepared the HN878-GFP recombinant strain. JP performed confocal imaging and prepared figures 3, 4 and S5. DS performed the histological analysis (Figure S4) and did the lung biopsy rejection. AL and BL performed lung surgery and obtained informed consent from the patients. LH isolated the SARS-CoV-2 strains. FCh processed PCSL samples by conventional TEM technique. JBG acquired the TEM images. FM, DB and NW provided scientific guidance for the project and critically reviewed the manuscript. All the co-authors read and approved the manuscript.

## Acknowledgments

We thank Sophie Guyetant (*Centre de ressources biologiques du CHRU de Tours*) and the *INRAE Val de Loire Partenariat* team for their assistance with the regulatory aspects of the study. We thank Fabienne ARCANGER and Juliette ROUSSEAU for their assistance with PCLS preparation for transmission electron microscopy (TEM). We also thank Marion Horta, who did her Master 1 internship with AR and assisted with PCLS preparation. We thank Roland Brosch (Institut Pasteur, Paris) who kindly provided the wildtype HN878 Mtb strain, and Sebastien Leclercq (ISP, INRAE, Nouzilly, France) who verified its genome sequence.

The development of these protocols was supported by the Microbiology Department of INRAE, INSERM, *Université de Tours* and Tours Regional and University Hospital. AR and SE received support from the French *Fédération de Recherche en Infectiologie* (FéRI). MSV holds a PhD fellowship funded jointly by the Loire Valley Region and the Microbiology Department of INRAE.

## Competing interests

None of the authors has any conflict of interest to declare.

## Supplementary materials and methods

### RNA extraction and gene expression analysis

Total RNA was extracted with the MagMAX™-96 Total RNA isolation kit (Thermo Fisher Scientific, ref. AM1840). The RNA was treated with DNase and mRNAs were then reverse transcribed with the iScript^TM^ Reverse Transcriptase mix (Biorad, ref 1708840) according to the manufacturer’s instructions. Primers (ordered from Eurogenetec; listed below) were validated on a serially diluted pool of cDNAs obtained from human lung biopsy samples and Vero cells infected with SARS-Cov-2, with a LightCycler® 480 Real-Time PCR System (Roche). Gene expression was then assessed with a BioMark HD (Fluidigm) in a 48 x 48 well IFC plate, according to the manufacturer’s instructions. The annealing temperature was 60 °C. Data were analyzed with Fluidigm RealTime PCR software to determine the cycle threshold (Ct) values. The expression of the SARS-CoV-2 *ip4* target region of *RdRp* was normalized against the mean levels of expression for four housekeeping genes (*ppid*, *gapdh*, *rps18* and *actb*) to obtain the ΔCt value. Data are expressed as relative amounts = 2^-ΔCt^).

### Primers used in this study

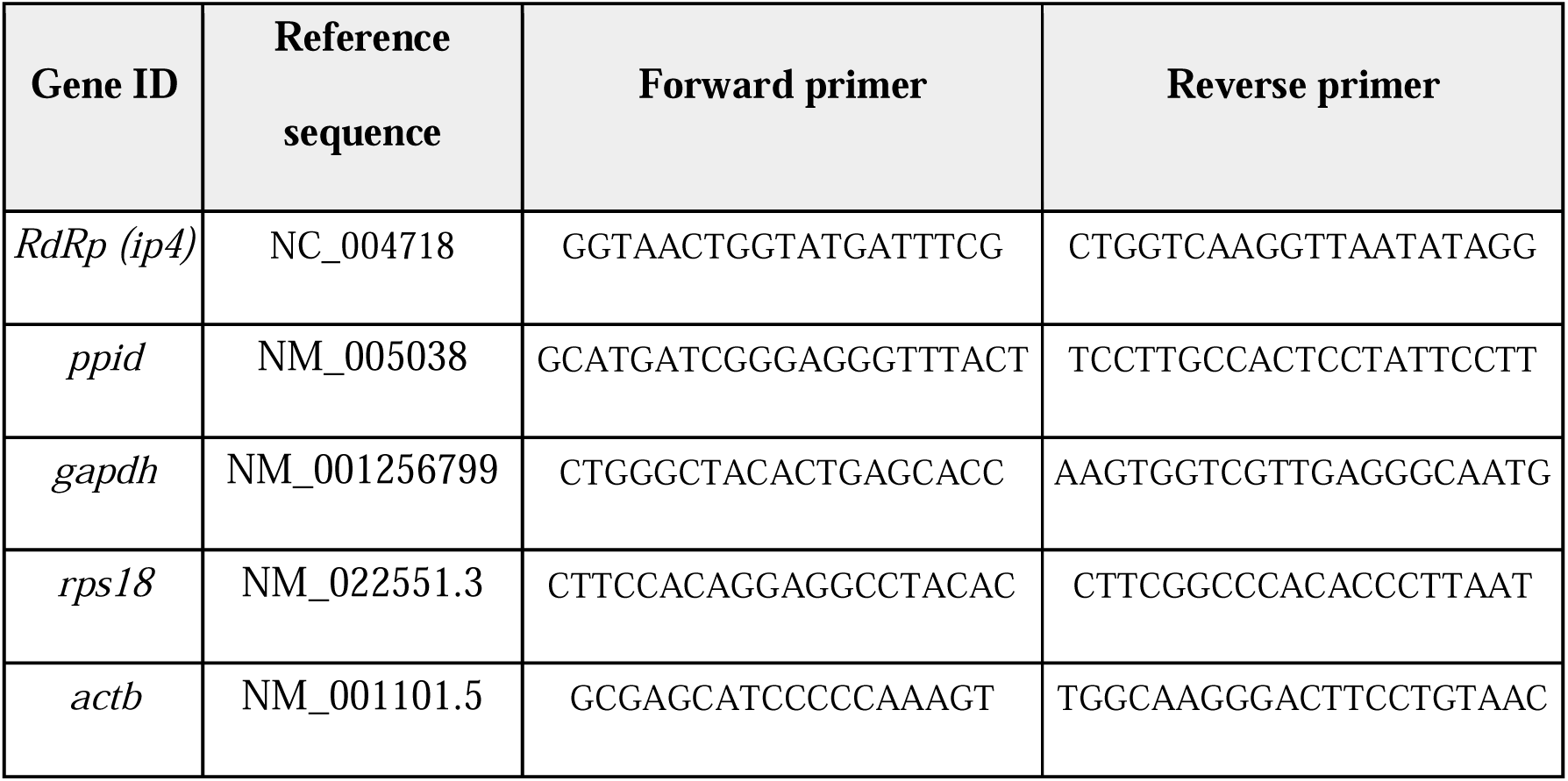

---

**Figure S1:**
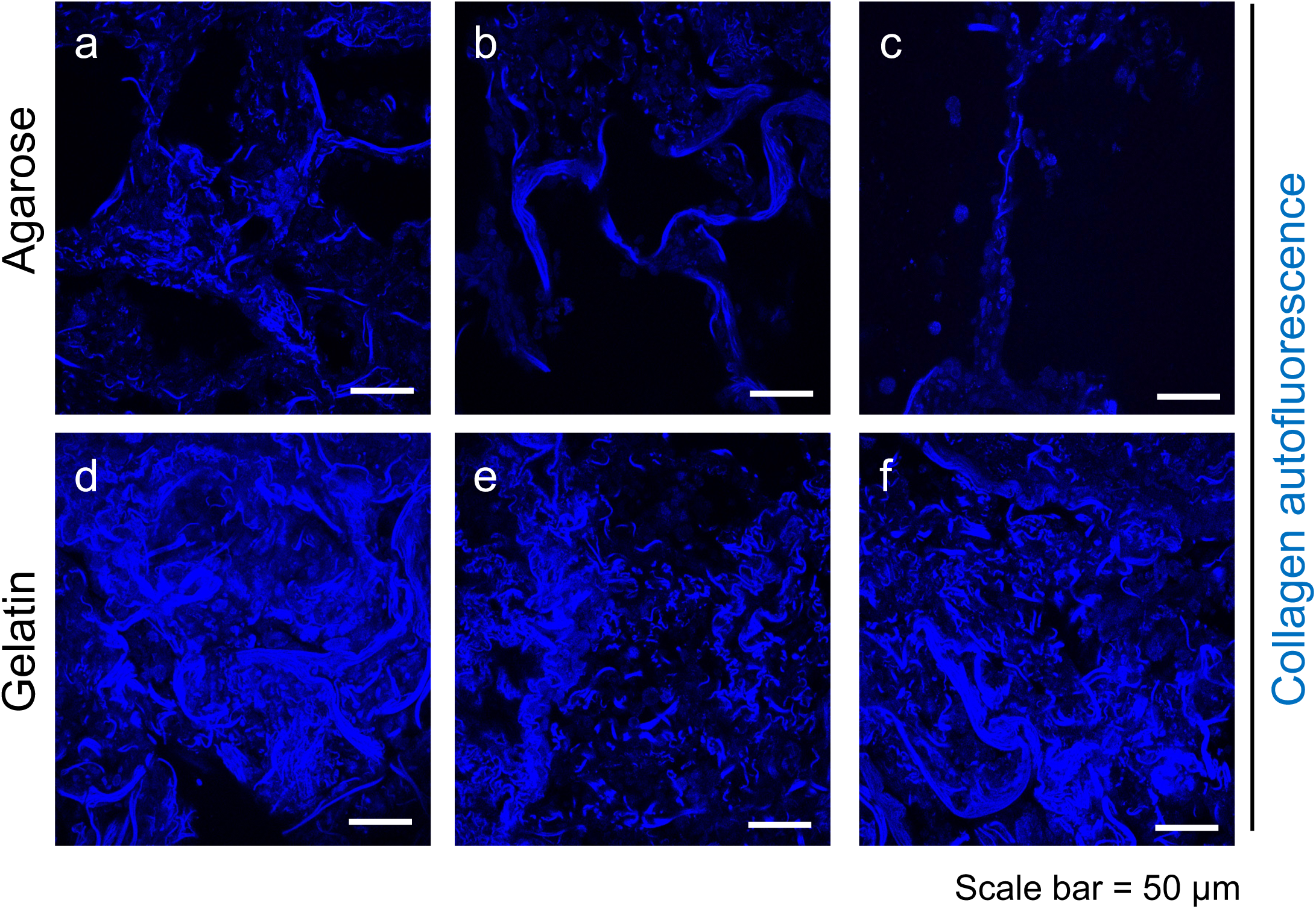
Comparison between low-melting-point agarose and gelatin. PCLS were obtained as described in figure 1 and filled with either low-melting point (LMP) agarose (a,b,c) or gelatin (d,e,f). The PCLS were fixed, labeled with Hoechst stain and observed with a confocal microscope. The alveolar sacs were filled and expanded with agarose but appeared to have collapsed with gelatin. Three representative images from one patient are shown (3 PCLS per filling agent, 2 independent experiments).

**Figure S2:**
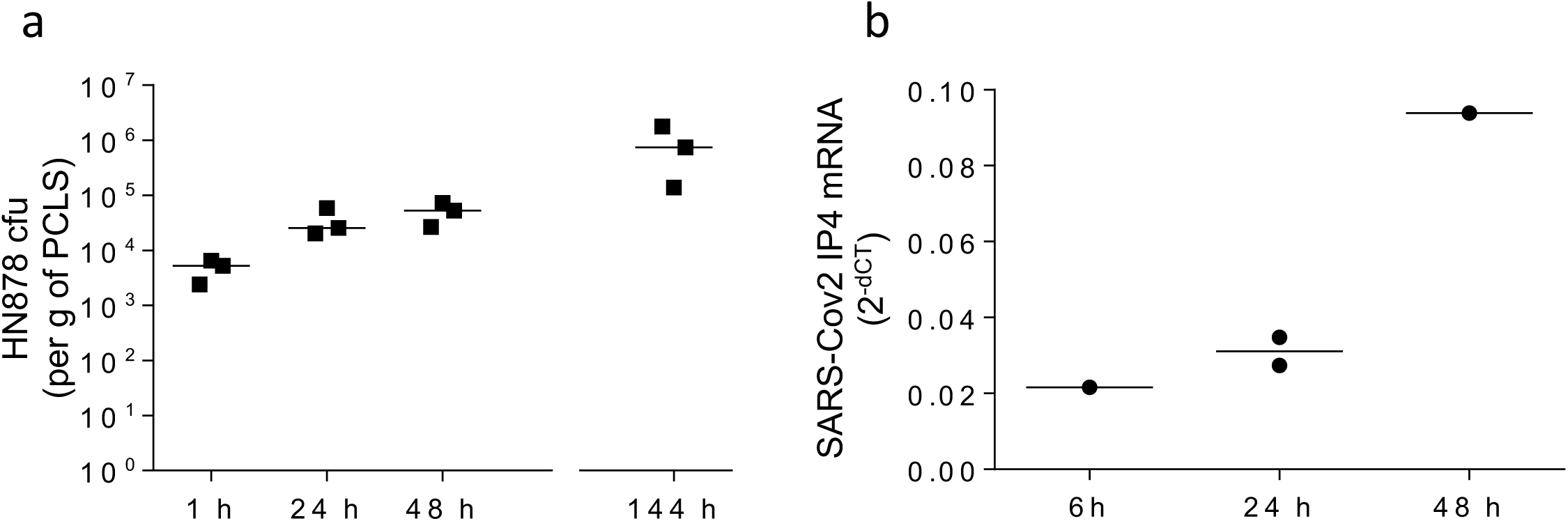
Mtb and SARS-CoV-2 replication in human PCLS. PCLS were obtained as described in figure 1. They were weighed on a precision scale and infected with a) 10^5^ CFU HN878 or b) 10^6^ PFU SARS-CoV-2 (Delta strain). The medium was replaced 24 h and 72 h post infection. a) At various time points post infection, PCLS were washed in DPBS and disrupted with a Precellys in Lysing Matrix D tubes containing DPBS supplemented with 0.1% Triton X-100. Tenfold serial dilutions were plated on 7H11 agar plates and CFU were counted two to three weeks later. CFU counts were normalized per g of PCLS. Median and individual data are shown (3 PCLS from the same patient). b) At different time points post infection, PCLS were washed in DPBS and disrupted with a Precellys in Lysing Matrix D tubes containing 800 µL Tri-reagent. RNA was extracted with the MagMax kit from Thermo Fisher Scientific in accordance with the manufacturer’s instructions. Reverse transcription was performed with iScript^TM^ kit from BioRad, and expression of the *ip4* target region of the *RdRp* gene was measured by RT-qPCR with normalization against the expression of four housekeeping genes (*ppid*, *gapdh*, *rps18* and *actb*). Individual data from one patient are shown.

**Figure S3:**
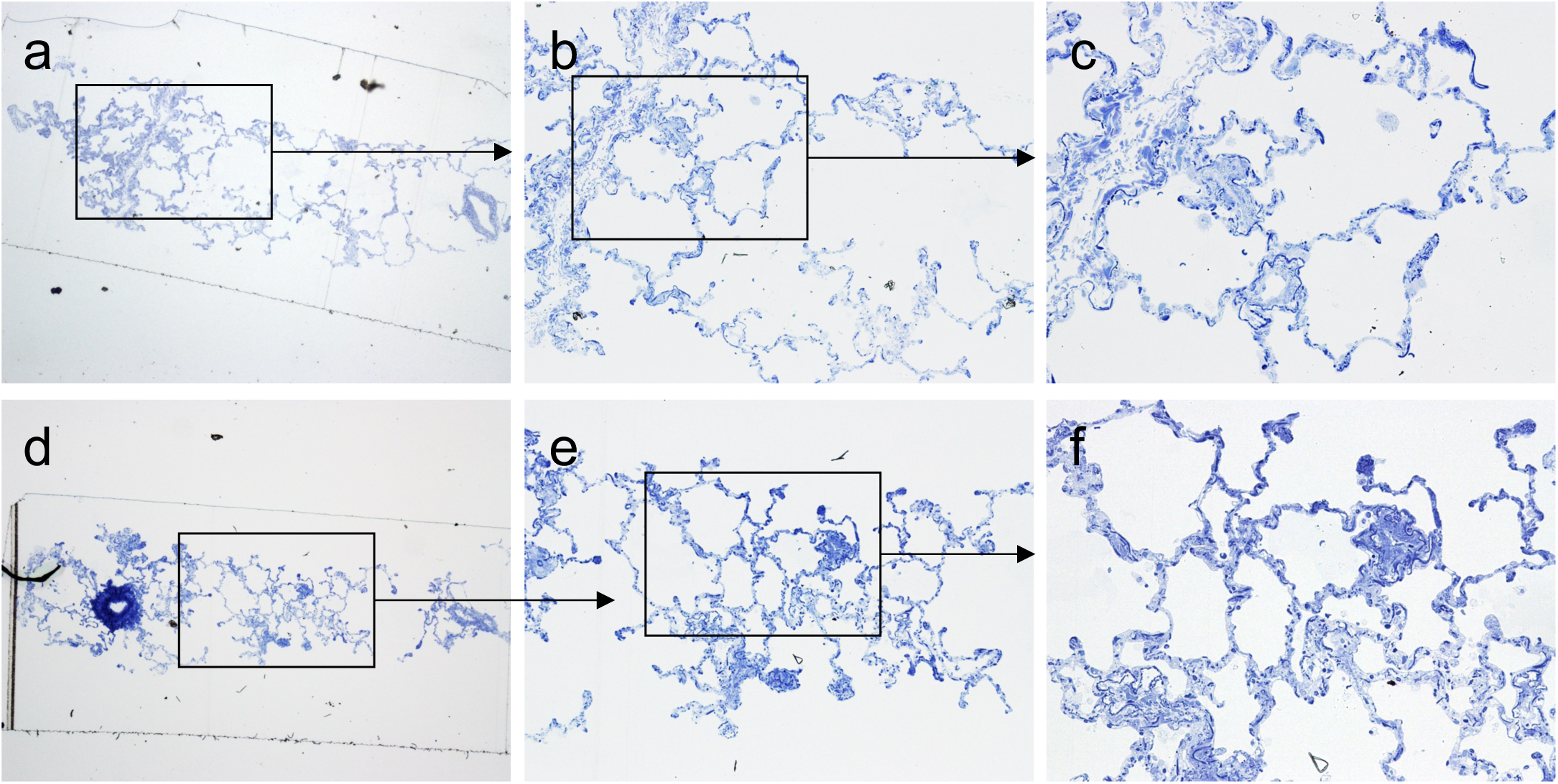
Toluidine blue staining on semi-thin section before the cutting of ultra-thin sections. PCLS were obtained as described in figure 1. PCLS were fixed in 1% glutaraldehyde, 4% paraformaldehyde and embedded in Epon resin to obtain a block. Semi-thin sections were cut from these blocks with an ultramicrotome, stained with toluidine blue, and observed under a light microscope. Observation of the semi-thin sections made it possible to localize areas of interest in the sample for the cutting of ultra-thin sections (Figure 5). The images (a,b,c) and (d,e,f) are progressive magnifications for two different patients.

**Figure S4:**
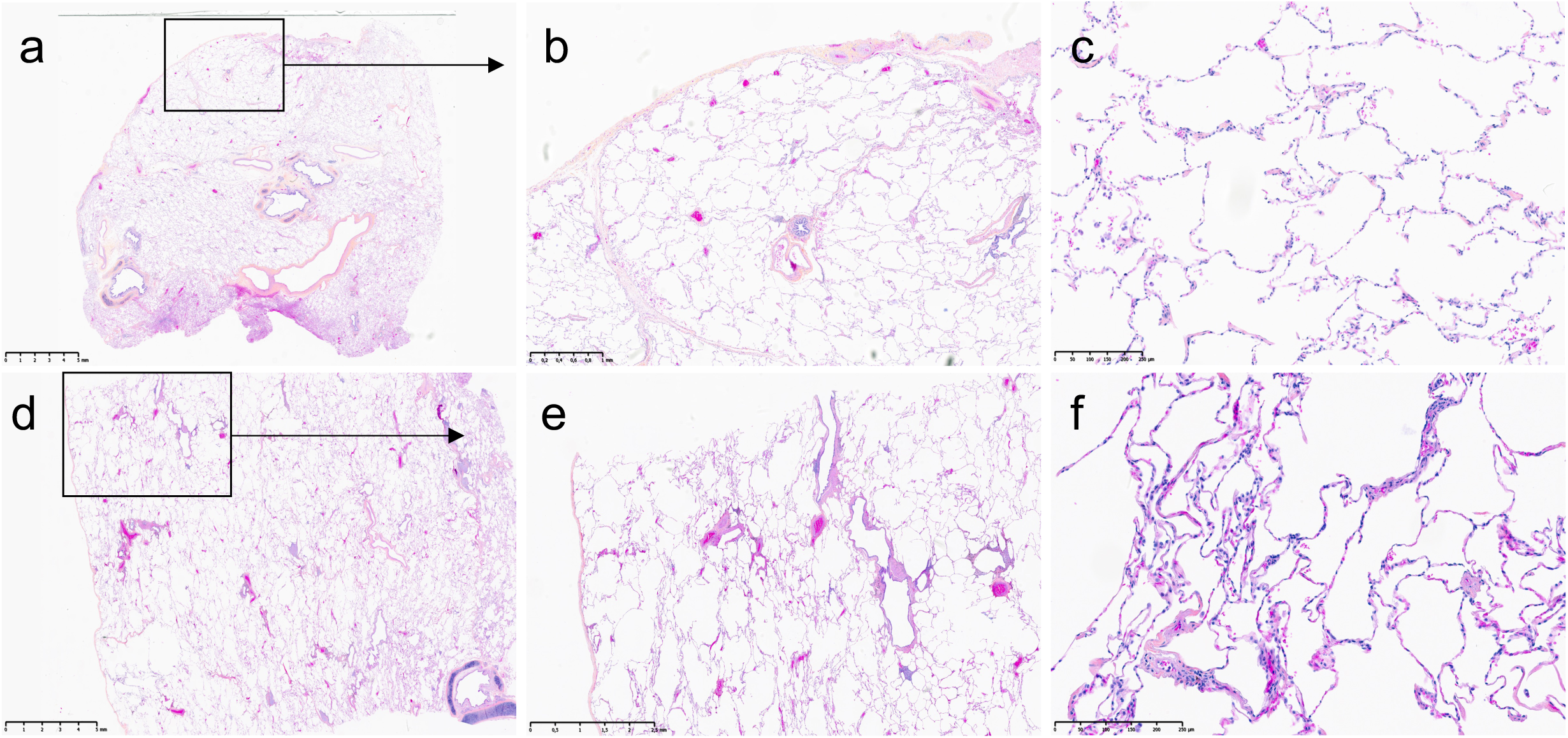
Histological evaluation of pulmonary tissue samples. During pathological diagnosis, a centimeter-sized tissue fragment was excised from the peripheral non-tumor area, embedded in paraffin and cut with a microtome. Sections were stained with hematoxylin/eosin to assess fibrosis, inflammation and epithelial degeneration. Lung parenchyma typically had a normal architecture with some subpleural dystrophic emphysematous lesions. The alveoli contained macrophages and the bronchovascular axes were normal. The images shown (a,b,c) and (d,e,f) are progressive magnifications from two different patients.

**Figure S5:**
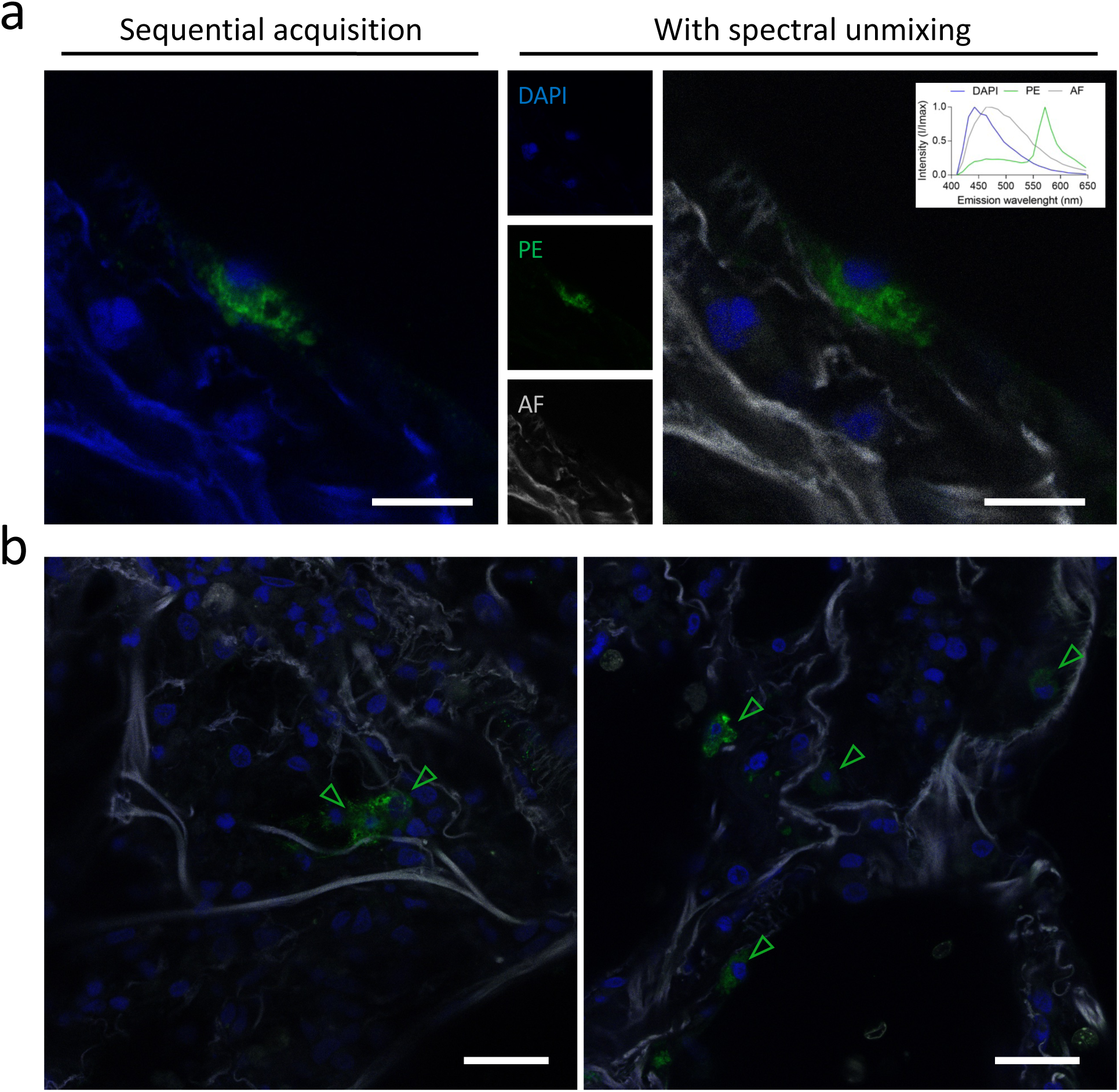
Spectral analysis of confocal images to remove autofluorescence background. (a) Imaging of a PCLS infected with SARS-CoV-2 by conventional sequential acquisition (left) or after spectral unmixing (right). Nuclei were counterstain with DAPI (blue) and infected cells were immunolabelled with anti-nucleocapsid antibody and detected with a secondary antibody coupled with phycoerythrin (PE; green). Autofluorescence is represented in gray following spectral unmixing. Scale bar[=[20 µm. (b) Spectral imaging of large areas of SARS-CoV-2-infected PCLS. Empty green arrowheads highlight infected cells. Scale bar = 25µm.

**Figure S6:**
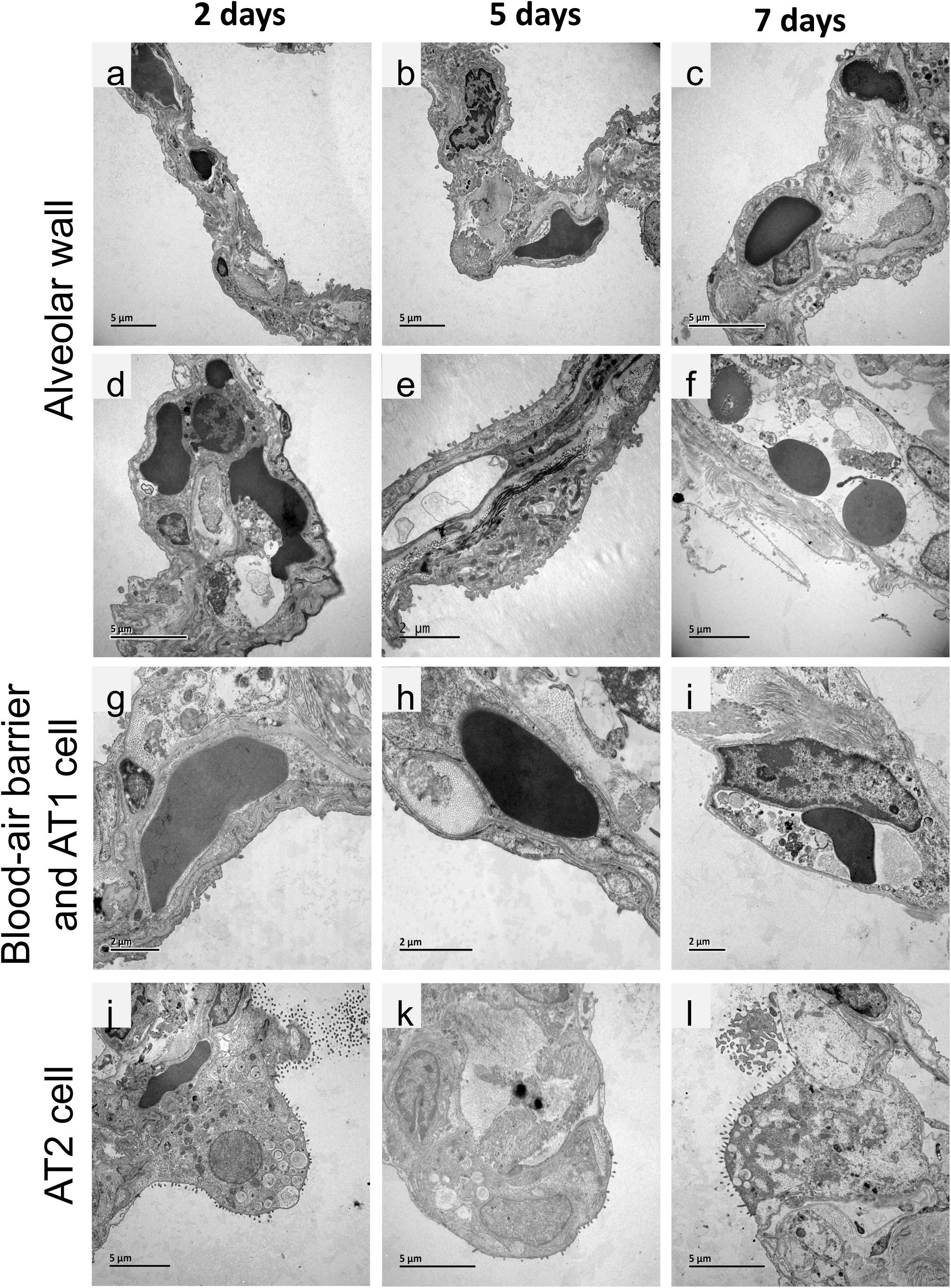
Preservation of alveolar wall ultrastructure over time, as visualized by TEM. PCLS were obtained as described in figure 1 and processed for TEM as described in figure 5. Non-infected PCLS were fixed at various times after obtaining the PCLS. The following structures are well preserved for up to five days: alveolar wall and connective tissue framework (a, b, d, e), blood-air barrier elements with endothelial cells and alveolar epithelial cells type I (g, h), and alveolar epithelial cells type II (j, k). At day 7, the ultrastructure of these components was significantly impaired (c, f, i, l). Representative images from one experiment of three performed are shown (1 patient per experiment; i.e., tested on 3 different patients).

**Figure S7:**
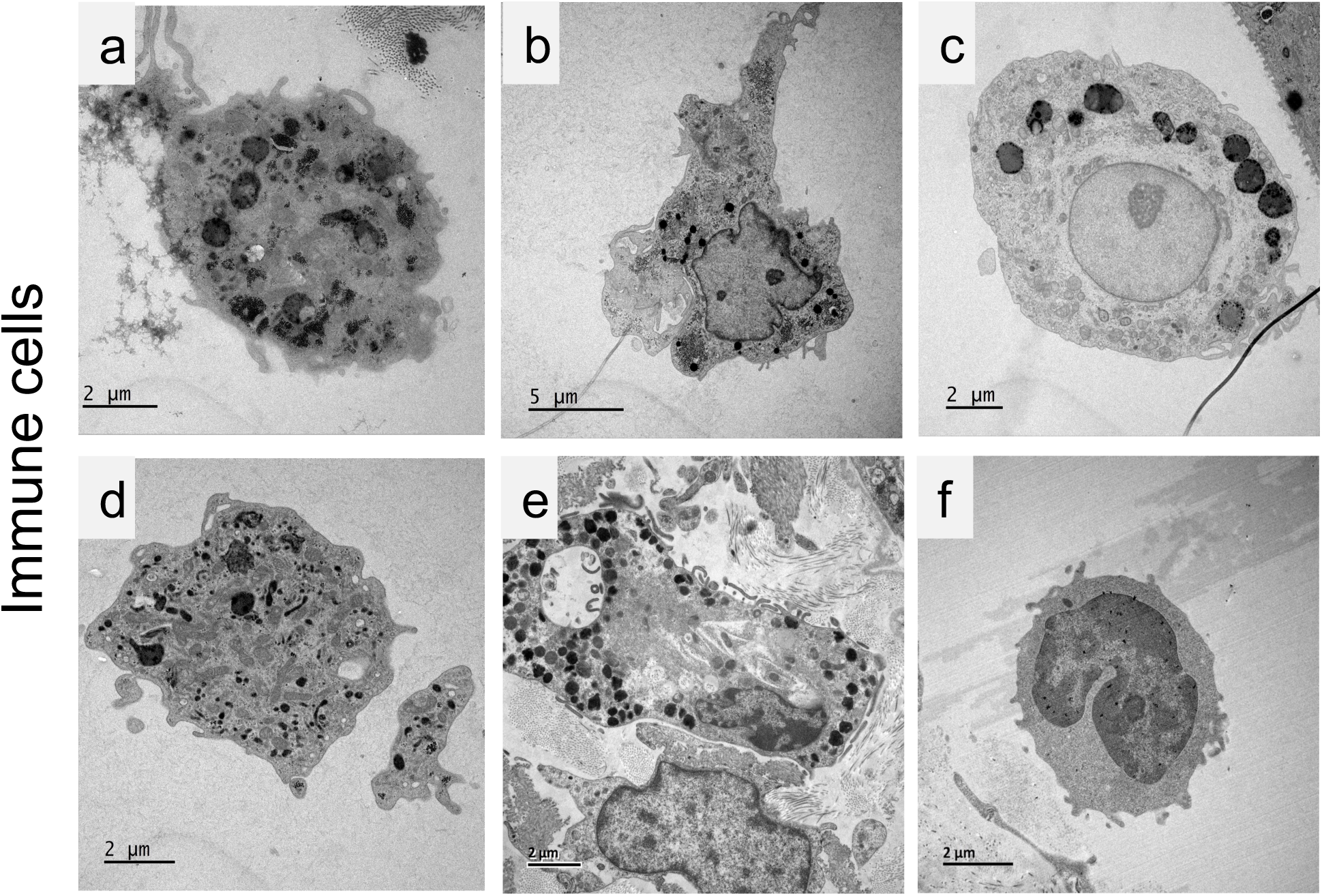
Preservation of alveolar immune cell ultrastructure visualized by TEM. PCLS were obtained as described in figure 1 and processed for TEM as described in figure 5. The following immune cells were observed, with well-preserved subcellular structures: alveolar macrophages (a to d), granulocytes (e), lymphocytes (f). Representative images from one experiment of three performed are shown (1 patient per experiment, i.e., tested on 3 different patients).

## Notes

### Competing Interest Statement

The authors have declared no competing interest.

